# Efficient and rapid fluorescent protein knock-in with universal donors in mammalian stem cells

**DOI:** 10.1101/2022.10.10.511307

**Authors:** Yu Shi, Nitya Kopparapu, Lauren Ohler, Daniel J. Dickinson

**Affiliations:** Department of Molecular Biosciences, the University of Texas at Austin

## Abstract

Fluorescent protein (FP) tagging is a foundational approach in cell biology because it allows observation of protein distribution, dynamics, and interaction with other proteins in living cells. However, the typical approach using overexpression of tagged proteins can perturb cell behavior and introduce localization artifacts. To preserve native expression, fluorescent proteins can be inserted directly into endogenous genes. This approach has been standard in yeast for decades, and more recently in invertebrate model organisms with the advent of CRISPR/Cas9. However, endogenous fluorescent protein tagging has not been widely used in mammalian cells due to inefficient homology-directed repair (HDR). Here, we describe a streamlined method for efficient and fast integration of FP tags into native loci via non-homologous end joining (NHEJ) in mouse embryonic stem cells. Our protocol minimizes cloning with universal donors, allows for N or C-terminal tagging of endogenous proteins, and requires less than 2 weeks from transfection to imaging, thereby improving the applicability of FP knock-in in mammalian cells.

## Introduction

Fluorescent protein (FP) knock-in enables endogenous tagging for protein visualization without overexpression artifacts^1^. A knock-in strategy allows an investigator to accurately observe and measure the dynamics of protein expression, localization, and interactions in live cells.

FP knock-in has been standard practice in yeast since the 1990s, because this organism can efficiently incorporate FP donors via homologous recombination^2,3^. More recently, FP knock-in has become widely adopted in *C. elegans*^4–7^ and *Drosophila*^8,9^ due to the advent of CRISPR/Cas9 technology^10^. When programmed by a single guide RNA (sgRNA), Cas9 introduces a targeted DNA double-strand break (DSB), which cells can repair by either homology-directed repair (HDR) or non-homologous end joining (NHEJ)^11^. HDR has been preferred due to its high fidelity^12–15^. However, HDR is only active at certain cell cycle stages^16^ and requires homology arms that match the target. As a result, HDR-based tagging is much less efficient^17,18^ and requires laborious cloning in mammalian cells.

To circumvent these constraints, NHEJ has been recently introduced for FP knock-in in mammalian cells^18–26^. One method, named CRISPR-assisted insertion tagging (CRISPaint) ^22^ is especially streamlined because it uses universal donor plasmids, such that the only cloning required is construction of the gene-specific sgRNA. The donor plasmid is introduced into cells via transfection, cleaved by Cas9 in parallel with the target gene, and integrated into the target gene in a non-sequence-specific manner by NHEJ. To allow the use of any gene-specific sgRNA while maintaining the correct reading frame, CRISPaint uses generic “frame selectors” to cleave the universal donor in one of the three possible reading frames^22^. Despite these advantages, to date CRISPaint has only been tested in a handful of cell lines. Additionally, only C-terminal insertion is feasible with the CRISPaint system in its present form, which has limited its application to genes whose protein products are tolerant of a C-terminal tag.

Here, we describe an improved approach, based on CRISPaint, that provides flexible and rapid FP tagging at either end of a gene in mammalian cells. Our approach is efficient, requires minimal cloning, and can produce functional endogenously tagged proteins expressed under the control of native regulatory elements. We tested and optimized this approach in mouse embryonic stem cells (mESCs). We successfully tagged 5/5 targets on the first attempt, and the time from transfection to imaging was as little as 2 weeks. Additionally, we have constructed a set of plasmids for multi-color tagging. Together, these advances will facilitate cell biological studies in mESCs and likely in other mammalian cells, and potentially offer a faster and easier route to quickly create knock-in mice.

## Results

### Adapting CRISPaint for C-terminal knock-ins in mESCs

CRISPaint ^22^ is a modular three-plasmid system which consists of an FP donor vector, a frame selector Cas9/sgRNA vector and a target Cas9/sgRNA vector (Figure S1a). The donor vector contains an mNeonGreen (mNG) fluorescent protein coding gene fused with a flexible linker, a self-cleaving T2A peptide, and a Puromycin resistance (PuroR) gene. The T2A peptide induces ribosomal skipping during translation, such that the mNG-fused protein of interest and the PuroR are transcribed as a single mRNA but produce two separate protein products^27,28^. A pre-made frame selector plasmid expresses Cas9 and an sgRNA that cleaves the linker region of the donor vector at one of three possible positions, chosen based on the target cleavage position to allow the linearized donor to incorporate into the cleaved gene in the correct reading frame. A second Cas9/sgRNA vector cleaves the gene of interest. Of note, the donor vector does not contain its own promoter, so the FP and PuroR sequences can only be expressed if the linearized donor is ligated to the cleaved gene in-frame and in the proper orientation (Figure S1a).

To date, the CRISPaint system has been tested in a handful of human cells^22,23,25^. In this study, we sought to expand the application of CRISPaint into mESCs which are a powerful and versatile model. mESCs are easy to maintain, can self-renew, and can divide indefinitely in defined growth medium^29^. Since they maintain naïve pluripotency, mESCs can develop into any type of somatic cells and can generate chimeric mice^30^. Moreover, mESCs are compatible with high-resolution microscopy because of low cellular autofluorescence.

To adapt CRISPaint for mESCs, we first replaced the CMV promoters, which were used to drive Cas9 expression, with EF1α promoters because the CMV promoter is inactive in mESCs^31^. As a first test, we attempted to tag the *Myh9* gene, encoding non-muscle Myosin IIA heavy chain, at the C-terminus with mNG (Figure 1a). *Myh9* was an appealing first target because it has a single annotated isoform and a known spatiotemporal localization to the cleavage furrow during mitosis, allowing us to use microscopy to assess whether the tagged protein localizes and functions normally in cell division^32^. We chose an sgRNA that cleaves near the C-terminus of the gene using CHOPCHOP^33^. The chosen sgRNA cuts 26 nucleotides (3N+2 nt, where N is the number of codons) away from the stop codon (Figure 1b). Since 2 nucleotides were lost at the Myosin cleavage site relative to the reading frame, we used frame selector +2 to cleave the donor vector, which should favor in-frame insertions by providing 2 extra nucleotides in linker region before the mNG coding sequence (Figure 1b). After transfecting and selecting Puromycin-resistant cells, we pooled putatively edited cells and tested for insertions using PCR with a forward primer upstream of the cut site and a reverse primer inside the mNG coding sequence (Figure 1c). We detected a specific 0.8 kb band in the pooled knock-in cells, indicating successful FP insertion (Figure 1c).

**Figure 1:**
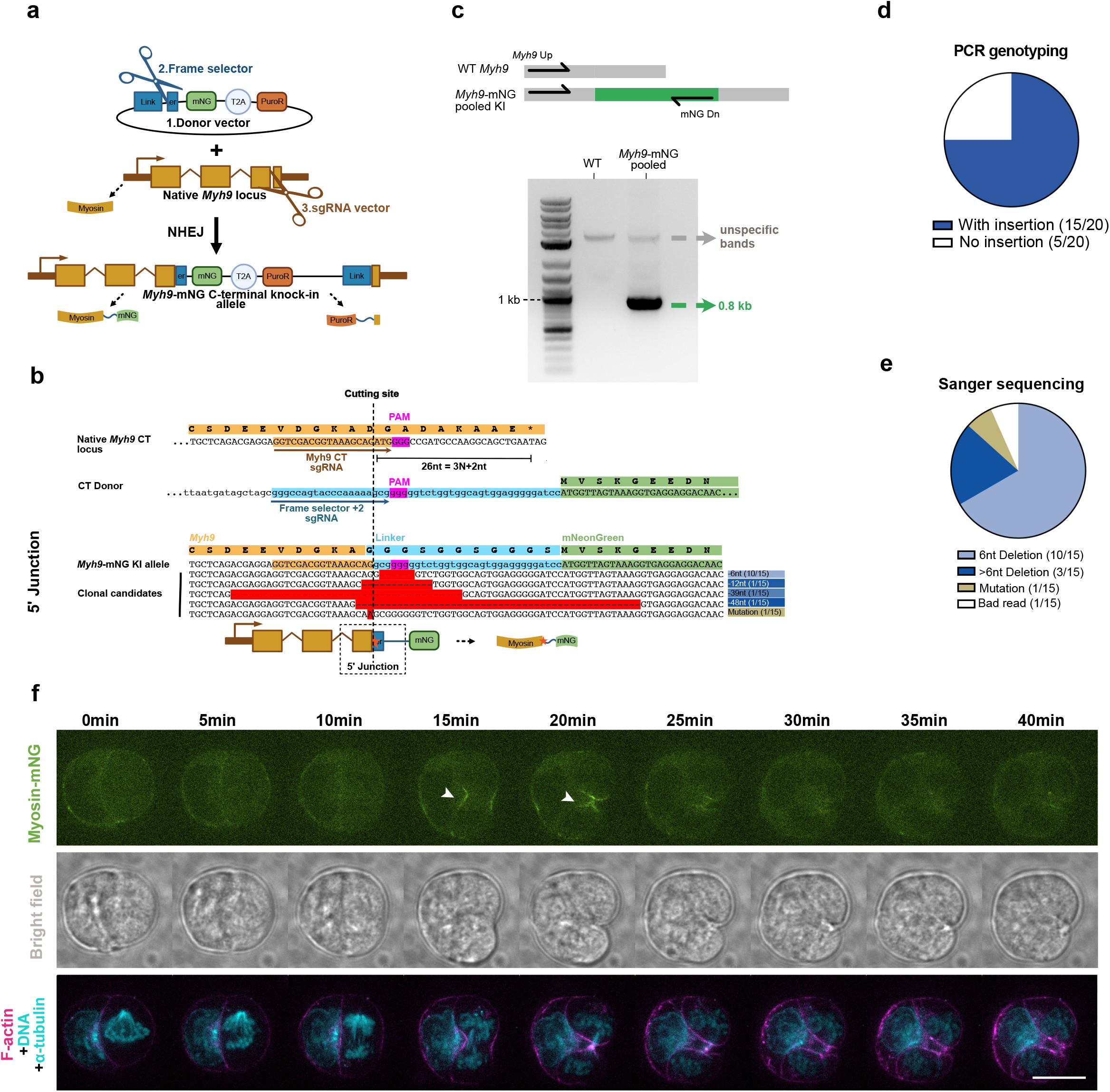
Adapting CRISPaint for C-terminal knock-ins in mESCs. **1a.**Illustration of C-terminal insertion via non-homologous end joining (NHEJ). The target sgRNA vector cuts the last exon of *Myh9* and the Frame selector cuts the linker region in the donor vector. The cleaved *Myh9* directly ligates with the linearized donor via NHEJ. T2A separates the knock-in gene into two protein products. **1b**. Sequence alignments showing the cleavage position of the target sgRNA in the target gene (top); the cleavage position of frame selector +2 in the donor vector (middle); and Sanger sequencing results in picked clonal cells (bottom). “-” in clonal sequences represents deletions. Highlighted nucleotides in clonal sequences indicate mutations. The red star in the cartoon indicates the locations of indels or mutations. **1c**. PCR genotyping of *Myh9*-mNG pooled cells. *Myh9* Up and mNG Dn primers amplified a specific band in *Myh9*-mNG pooled cells compared with WT cells, indicating a successful insertion. **1d**. Pie chart showing correct insertion efficiency in picked clonal cells. **1e**. Pie chart summarizing frequency of different insertion results in picked clonal cells. Bad read means an inconclusive Sanger sequencing result. **1f**. Timelapse live imaging of myosin-mNG during cell division. White arrowhead in the myosin-mNG channel highlights myosin-mNG accumulation at the cleavage furrow in dividing cells, indicating that the tagged protein is functional. Scale bar represents 10 µm. Abbreviations: C-terminal (CT), mNeonGreen (mNG), Puromycin resistance (PuroR), the T2A peptide (T2A), Wildtype (WT), Knock-in (KI), nucleotides (nt), 3N+2nt where N represents the codon number.

We next proceeded to isolate single knock-in clones from the pool of putatively edited cells. PCR genotyping showed that 15/20 clones had an integration of mNG, suggesting a high insertion efficiency (Figure 1d, S1b). Sanger sequencing revealed one clone with a single point mutation at the junction between *Myh9* and mNG, and 14 clones with small indels at the junction. The most frequent deletion, removing six nucleotides (encoding two glycine residues) from the linker region of the universal donor vector, was observed in 10 independent clones (Figure 1b,e). The presence of these indels was surprising because in human cells, CRISPaint resulted in 61.5–86.7% seamless insertions when using the correct frame selector^22^. This difference may be due to differences in NHEJ mechanisms in mESCs compared to human cells.

There are at least two distinct NHEJ pathways, termed alternative NHEJ (aNHEJ) and classical NHEJ (cNHEJ), which produce indels at different frequencies and may have different relative activity levels in different cells^34,35^. Nevertheless, all the knock-in clones we isolated had deletions whose length was a multiple of three, resulting in an in-frame fusion of mNG to the C-terminus of Myosin, as expected (Figure 1b,e). The loss of six nucleotides (two amino acids) from the linker region of the donor vector is expected to be inconsequential for protein function. Indeed, live imaging confirmed that Myosin-mNG accumulated at the cleavage furrow in dividing cells, indicating the tagged protein is functional (Figure 1f, movie 1). In summary, we successfully adapted CRISPaint for use in mESCs, and we achieved efficient and functional tagging at the C-terminus of Myosin IIA despite small in-frame deletions at the junctions.

A potential concern when using this approach for gene tagging is that bacterial sequences, present in the donor plasmid backbone, could interfere with gene expression by disrupting the 3’UTR or other regulatory elements. This issue can be avoided by generating “minicircle” donors from which the bacterial sequences have been removed^22^. To confirm that the minicircle strategy is applicable in mESCs, we cloned a C-terminal parent vector in which attB-attP recombination sites flank the Linker–mNG–T2A–PuroR cassette (Figure S1c). In this way, attB/attP recombination circularizes the donor sequences, and the bacterial backbone is eliminated due to cleavage at its 32 I-Scels restriction sites. We induced minicircle production in an engineered *E. coli* strain^36^ and purified the minicircle donor (Figure S1c-d). We note that although we initially tested published protocols for minicircle production^36^, we found that extensive further optimization was needed to eliminate bacterial DNA contamination and obtain sufficient yields for transfection (unpublished observations; see Supplemental Protocol for details). We transfected the C-terminal mNG minicircle donor into mESCs along with the same *Myh9* sgRNA and frame selector as before (Figure S1e). PCR genotyping of pooled edited cells demonstrated that the C-terminal minicircle donor was successfully inserted into the *Myh9* locus (Figure S1f).

### A universal donor for N-terminal tagging

The current version of CRISPaint is limited to C-terminal tagging, but some proteins have functional domains or motifs at their C-termini which may be sterically hindered by fluorescent protein tagging^37^. Therefore, we sought to develop a similar approach for introducing N-terminal tags. There are several important considerations for N-terminal tagging. First, to ensure gene expression, the insertion must be in-frame at both junctions. Second, no potential stop codons should exist inside the inserted sequence. This poses a challenge if the donor DNA contains the entire bacterial plasmid backbone; therefore, a minicircle donor is required. Third, the order of mNG and PuroR is important; the mNG needs to be adjacent to the gene of interest so that the mNG tag is not removed by T2A peptide cleavage.

We designed a novel N-terminal parent vector by making several changes to the C-terminal vector design (Figure S2a-b). To fuse mNG with the N-terminus of a gene, we rearranged the position of the mNG and PuroR related to T2A peptide. To avoid putative stop codons, we flanked the attL sequence, which was left over from removal of the bacterial backbone during minicircle donor production, with splicing signals to create an artificial intron^38^. To facilitate in-frame insertion, we redesigned the linker to have a length that is a multiple of three. We produced new frame selector sgRNA vectors for the N-terminal minicircle donor, which can result in a correct reading frame on both 5’ and 3’ junctions.

We tested our N-terminal minicircle donor strategy by tagging *Myh9* at the N-terminus (Figure 2a). For N-terminal knock-in, we selected an sgRNA that cleaves just before the start codon. Since this cut site is 3N+2nt away from the stop codon, we used frame selector -2 to favor in-frame tagging (Figure 2b). After co-transfection and drug selection, PCR genotyping indicated successful insertion in pooled edited cells (Figure 2c). PCR genotyping of clonal cells showed that 9/12 clones had correct insertions with a specific 1.8kb band (Figure 2d, S2c). Two of the incorrect clones had multiple copies of the minicircle donor inserted head-to-tail into the cut site (Figure S2c, S2f); these clones were successfully edited but are not usable for imaging due to the presence of an additional copy of mNG that is not fused to Myosin.

**Figure 2:**
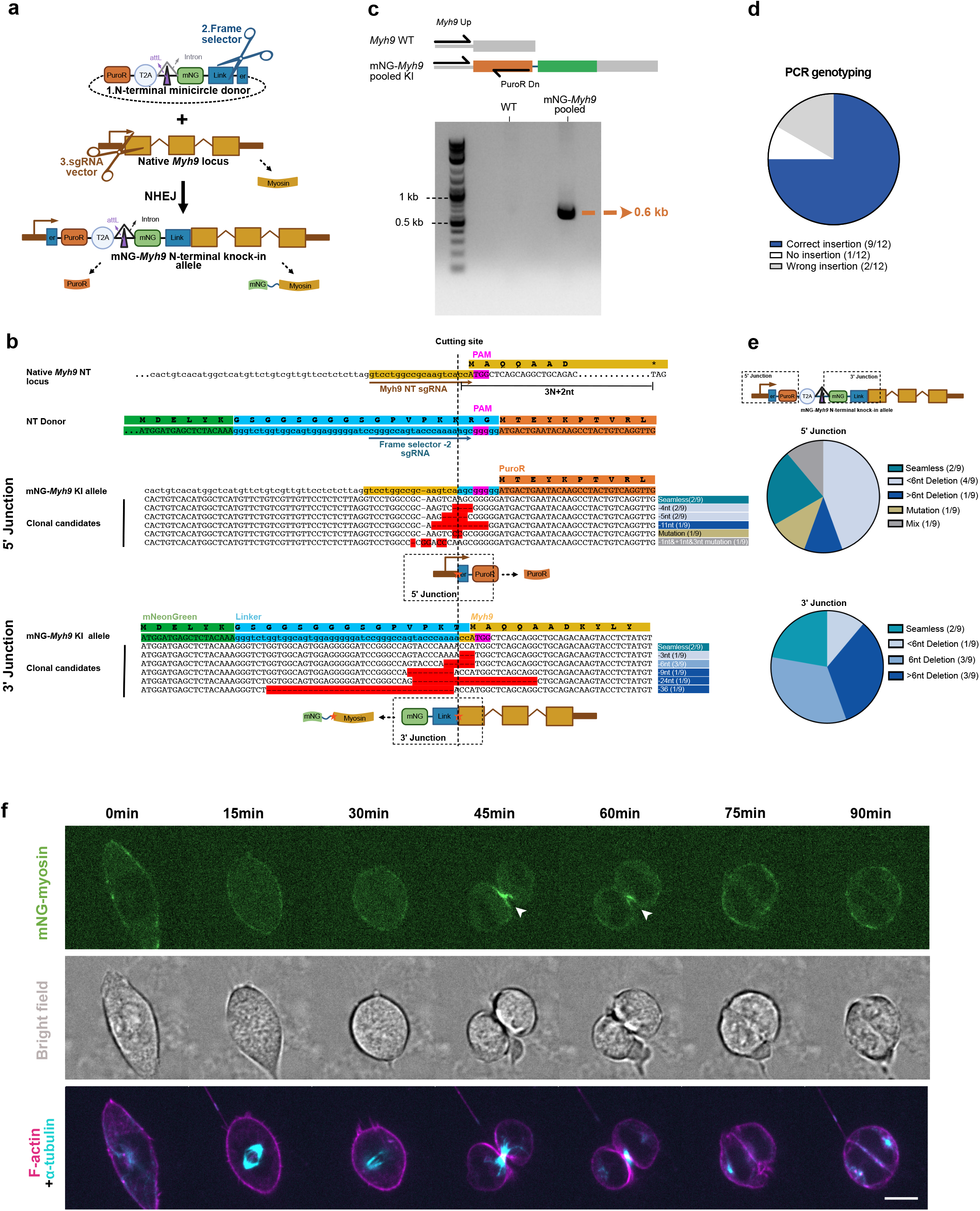
Efficient knock-in at N-terminus of myosin. **2a.**Illustration of N-terminal insertion via non-homologous end joining (NHEJ). The target sgRNA vector cuts the 5’ UTR of *Myh9* and the frame selector cuts the linker region of the N-terminal minicircle vector. The cleaved *Myh9* directly ligates with the linearized donor via NHEJ. T2A separates the knock-in gene into two protein products. **2b**. Sequence alignments showing the cleavage position of the target sgRNA in the target gene (top); the cleavage position of frame selector -2 in the donor vector (middle); and Sanger sequencing results for both the 5’ and 3’ junctions in picked clonal cells (bottom). “-” in clonal sequences represents deletions. “-” in the reference sequence represents insertions. Highlighted nucleotides in clonal sequences indicate mutations. The red star in the cartoon indicates the locations of indels or mutations. **2c**. PCR genotyping of mNG-*Myh9* pooled cells. *Myh9* Up PuroR Dn primers amplified a specific band in mNG-*Myh9* pooled cells compared with WT cells, indicating a successful insertion. **2d**. Pie chart showing correct insertion efficiency in picked clonal cells. **2e**. Pie charts summarizing frequency of different insertion results in both junctions. Mix means a mixture of indels or/and mutations. **2f**. Timelapse live imaging of mNG-myosin during cell division. White arrowhead in the mNG-myosin channel highlights mNG-myosin accumulation at the cleavage furrow in dividing cells, indicating that the tagged protein is functional. Scale bar represents 10 µm. Abbreviations: N-terminal (NT), mNeonGreen (mNG), Puromycin resistance gene (PuroR), the T2A peptide (T2A), Wildtype (WT), Knock-in (KI), nucleotides (nt), 3N+2nt where N represents the codon number.

Sanger sequencing of the 9 clonal cell lines that had correct insertions identified one insertion with a seamless 5’ junction, one with a seamless 3’ junction, and one that was seamless at both junctions (Figure 2b,e and S2d). Thus, it is possible to obtain clones with seamless insertion junctions in mESCs, perhaps with variable frequency at different loci. As expected, among the clones that had indels, all clones maintained the correct reading frame at the junction where the 3’ end of the donor was fused to Myosin (Figure 2b,e). However, at the 5’ end of the donor, which is outside the coding region, 6/9 clones had deletions whose lengths were not a multiple of 3. This suggests that the length of indels produced by NHEJ in mESCs is essentially random, but that drug selection can effectively enforce an in-frame fusion between the tag and the protein of interest. Timelapse live imaging suggested that the N-terminally tagged mNG-Myosin is functional and localized normally to cleavage furrows in cells undergoing cytokinesis (Figure 2f, movie 2). The fluorescence signal was consistent among different knock-in clones (Figure S2e), suggesting that the small indels we observed did not affect Myosin expression or localization.

Since all the 12 mNG-Myosin clones we isolated were mono-allelic (i.e., only one copy of *Myh9* was tagged), we wondered if the sgRNA we chose was inefficient and had failed to cleave the other allele. Thus, we also sequenced the non-tagged allele in 9 clones which had a correct insertion on one allele. Sanger sequencing revealed indels or mutations at the non-tagged allele in 9/9 clones (Figure S1d). Thus, the sgRNA vector was efficient enough to cut both alleles, but in most cases only one allele was repaired via integration of the donor. In summary, we designed and produced N-terminal minicircle donors that can achieve efficient N-terminal endogenous tagging via NHEJ.

### NHEJ-based knock-in is rapid, robust, and applicable to many genes

To demonstrate the generality of this approach, we targeted more genes: Tpx2, a key mitotic spindle assembly factor; β-catenin, an adherens junction protein; and Zo1, a tight junction protein. We had initially targeted *Myh9* because this gene has only a single annotated isoform; however, these additional targets each have more than one isoform due to alternative splicing (Figure S3a-c). In this case, one key consideration for FP knock-in is which region of genomic DNA to tag. We chose to target the principal isoform as identified by APPRIS^39^, which flags the principal isoform based on protein structure, expression, and conserved function (Figure S3a-c). Based on the gene structure, we targeted the C-terminus of each gene, and we used C-terminal minicircle donors (Figure S1d) to avoid the possibility of disrupting each gene’s 3’UTR.

We readily obtained knock-in cells with tagged Tpx2, β-catenin and Zo1 on the first attempt. Insertion was verified by PCR using pooled edited cells after transfection and selection (Figure S3d-f). Fluorescence imaging of all three sets of pooled cells showed that most cells were fluorescent, suggesting that knock-in efficiency was high (Figure 3a-c). The pooled cells were immediately used for live imaging, which was possible within 7 to14 days after transfection. All three tagged proteins exhibited the expected localization. Tpx2-mNG cells showed faint nuclear fluorescence in non-dividing cells, but bright mitotic spindle localization in dividing cells (Figure 3a). Timelapse live imaging of Tpx2-mNG cells revealed dynamic localization of Tpx2 during mitosis (Figure 3d, movie 3). β-catenin-mNG cells showed cortical accumulation and brighter signal at cell-cell junctions (Figure 3d). Timelapse live imaging of β-catenin-mNG cells revealed *de novo* adherens junction assembly following cell division (Figure 3e, movie 4). Finally, Zo1-mNG cells displayed puncta on the cell cortex, which were concentrated at cell-cell contacts (Figure 3c).

**Figure 3:**
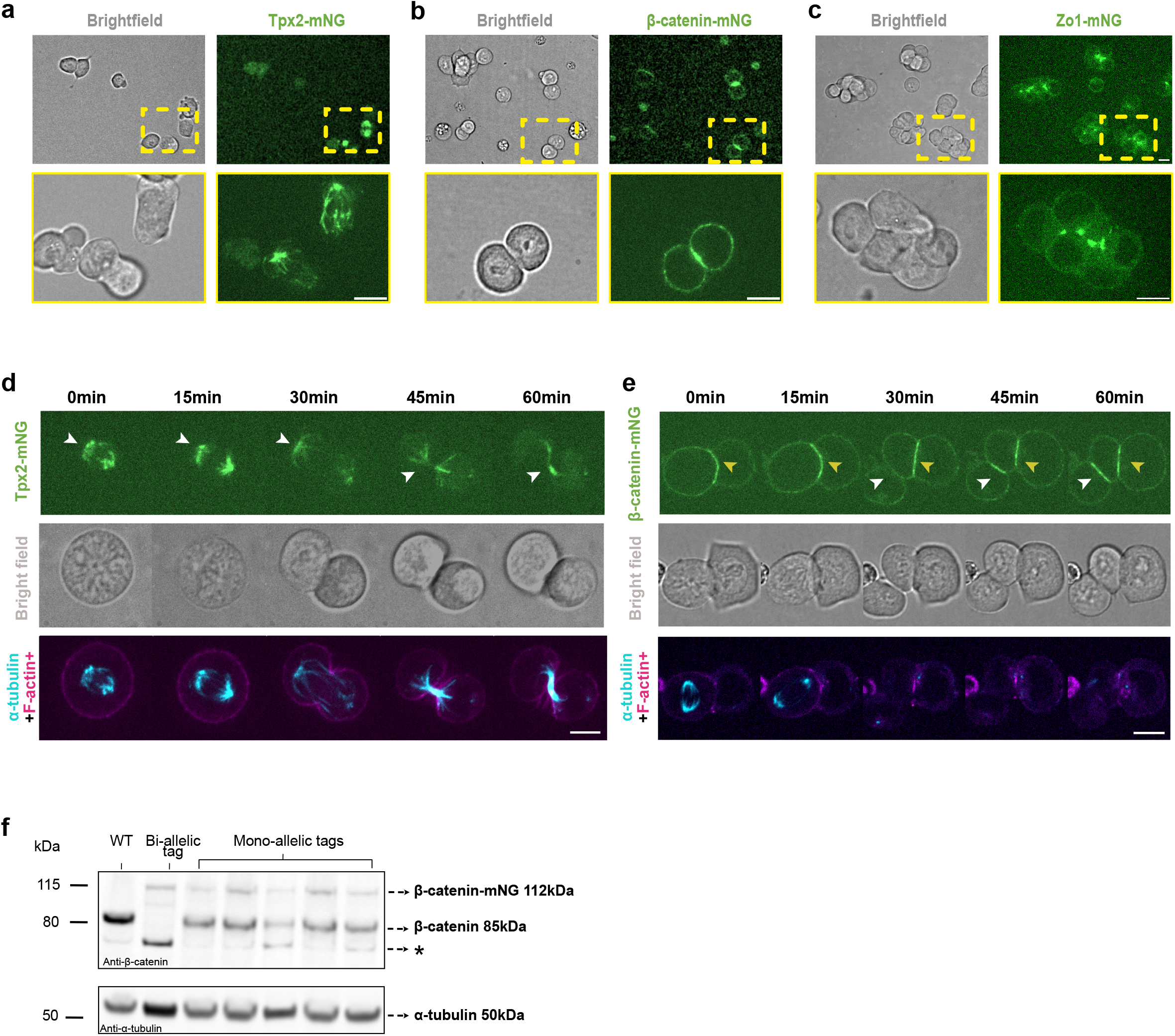
Tagging of additional genes and imaging of pooled edited cells. **3a-c.** Imaging results showing expression and spatial distribution of Tpx2-mNG, β-catenin-mNG, Zo1-mNG in pooled knock-in cells. **3d-e**. Timelapse live imaging of Tpx2-mNG and β-catenin-mNG during cell division. White arrowhead in the Tpx2-mNG channel highlights localization at the mitotic spindle during mitosis. White arrowhead in the β-catenin-mNG channel highlights *de novo* adherens junction assembly following cell division. Yellow arrowhead indicates a pre-existing adherens junction. Scale bars represent 10 µm. **3f**. Western blotting β-catenin-mNG clonal cells with mono-allelic or bi-allelic tag insertions. Asterisk indicates an unidentified band that may be non-specific or may represent a minor isoform of β-catenin.

To further validate the expected localization of mNG-Myosin IIA, β-catenin-mNG and Zo1-mNG, we examined the localization of these proteins in 3D culture by embedding cells in matrigel (Figure 4). Naïve mESCs grow in 3D culture as disorganized clusters, but when mESCs exit naïve pluripotency in 3D culture, they establish epithelial apicobasal polarity, forming a sphere with an open lumen at the center^40,41^. In polarized mESCs, β-catenin-mNG localized at cell-cell contacts and was enriched at the apical adherens junction. Zo1-mNG localized to the apical edge of cell-cell contacts, consistent with the expected position of tight junctions in a polarized epithelium^42^. Interestingly, mNG-Myosin showed both apical and basal accumulation in polarized mESCs. The apical enrichment was not unexpected since epithelia often enrich Myosin apically^42^ but the basal localization is novel, to our knowledge, and suggests that Myosin might play a role in organizing the cells into a sphere. As expected, none of these proteins showed a polarized localization in naïve mESCs. Of note, the imaging experiments using β-catenin-mNG and Zo1-mNG was performed using pooled puromycin-resistant cells 10 days after transfection, further demonstrating that our protocol can allow tagging and imaging of endogenous proteins in less than two weeks. In summary, live imaging of our knock-in cells in 3D culture further validates that these tagged proteins are functional and shows that endogenous tagging can reveal changes in protein localization and cell organization that are a consequence of cell fate transitions.

**Figure 4:**
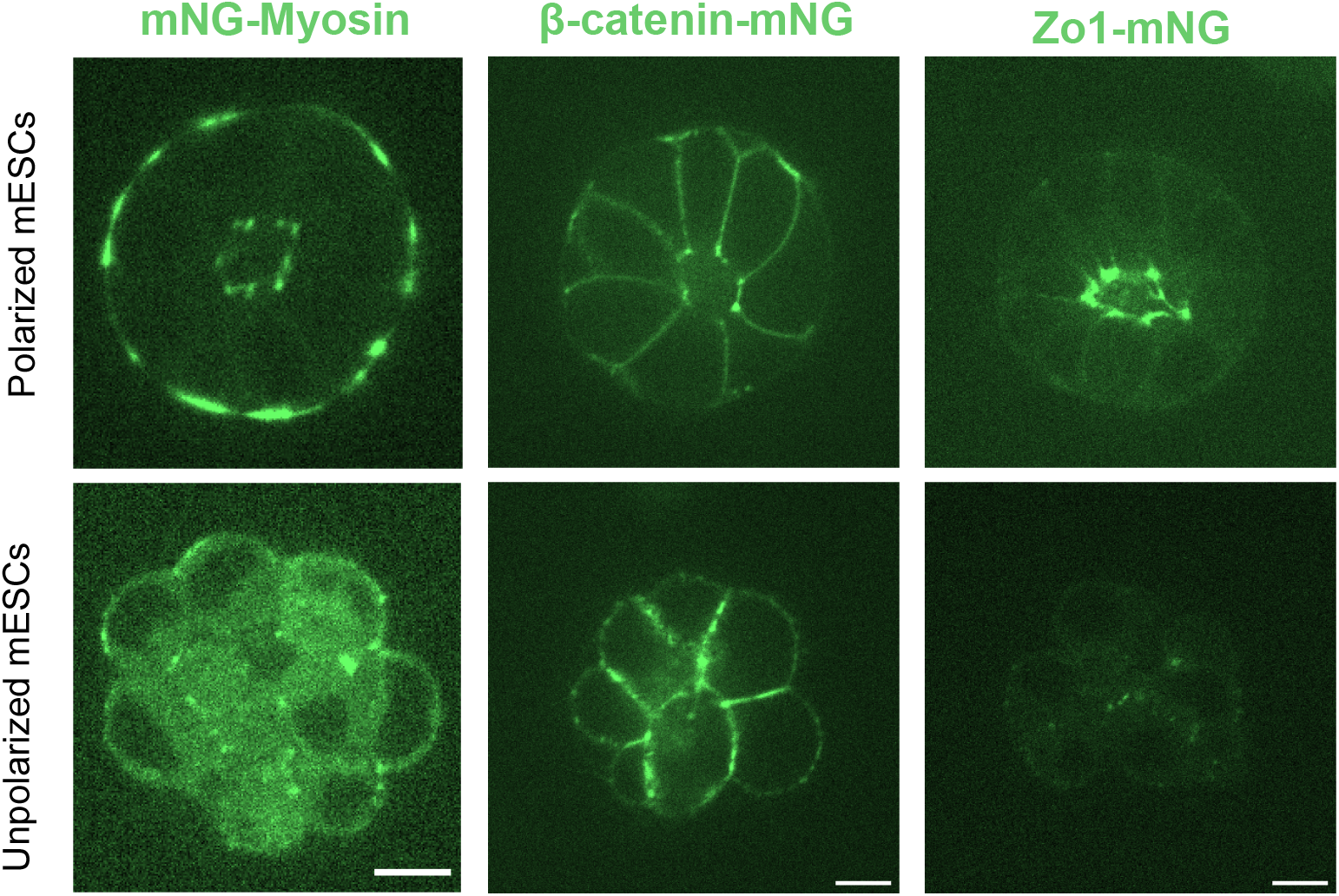
Epithelial polarization in knock-in mESCs in 3D culture. Images of mNG-Myosin, β-catenin-mNG, Zo1-mNG in living mESCs in 3D spheroid culture. Non-polarized mESCs were cultured in 2i/LIF, and polarized mESCs were grown without 2i/LIF. Scale bars represent 10 µm.

We also picked monoclonal cells carrying tagged Tpx2 and β-catenin. As expected, the insertion efficiency was high: 15/18 of Tpx2 clones (Figure S4a-b) and 33/36 β-catenin clones (Figure S5a-b) had correct insertions. Sanger sequencing revealed a few seamless insertion junctions along with small insertions, deletions, and mixtures of those (Figure S4c-d, S5c-d), but all tags were inserted in-frame. Additionally, we isolated two β-catenin-mNG clones that had both alleles tagged, demonstrating that bi-allelic tagging is possible (Figure S5a). Western blotting verified the expected pattern of β-catenin expression in cells carrying mono-allelic and bi-allelic tags: the cells with the bi-allelic tags lacked untagged β-catenin protein while the cells with mono-allelic tags expressed both the tagged and untagged forms (Figure 3f). Interestingly, although the tagged copy of β-catenin-mNG was expressed and localized normally, the tagged protein appeared was present at somewhat lower levels than the wild-type (untagged) protein (Figure 3f). This was not due to incomplete cleavage of the T2A peptide, since western blotting showed no evidence for a β-catenin-mNG-PuroR fusion (Figure S5e). Although the reason for lower relative expression of β-catenin-mNG relative to untagged β-catenin is unclear, the C-terminal tail of β-catenin has been shown to influence the binding affinity of β-catenin for the destruction complex components APC and Axin^43^, so it is conceivable that the presence of a C-terminal tag could change the stability of the β-catenin-mNG protein. This reduction in levels of β-catenin- mNG relative to untagged β-catenin did not cause any evident phenotypes, nor prevent us from observing β-catenin-mNG localization (Figures 3-4).

In our earlier experiments targeting *Myh9*, we exclusively isolated clones that had one tagged allele; the other allele was cleaved by the target sgRNA but not tagged. This suggested that ligation of the donor was a limiting factor, and we wondered whether using a higher concentration of donor would improve the efficiency of FP knock-in and bi-allelic tagging. Thus, we tested transfections with different [donor vector] : [frame selector] : [target sgRNA] plasmid moles ratios, ranging from 5:1:1 to 50:1:1. Regardless of the transfection ratio, PCR results confirmed an insertion in pooled cells in every experiment (Figure S3d-f). Thus, all ratios can generate knock-ins with some efficiency. We quantified the frequency of FP+ cells in each experiment and found that the fraction of FP+ cells ranged from 36% to 87%, any of which would be sufficient to allow imaging experiments in pooled cells without the need for clonal selection (Figure S3g). When targeting Tpx2, we observed a trend of increasing efficiency with higher transfection ratio of donor, but no clear pattern was observed when targeting β-catenin (Figure S3g). These data suggest that in some cases, using more donor might favor integration, but tagging can succeed regardless of the amount of donor within the range we tested.

Overall, we conclude that by combining NHEJ and drug selection, endogenous tagging in mESCs is generally very efficient. Although most cell clones only had one allele edited, bi-allelic edits can be isolated by picking single clones. Due to high insertion efficiency, pooled edited cells can be imaged less than two weeks after transfection.

### Additional vectors for dual-color knock-in by sequential transfections

To enable simultaneous labeling of two proteins with different tags in the same cell line, we constructed donor vectors for HaloTag insertion at either terminus of a protein (Figure 5a). HaloTag is a self-labeling enzymatic tag that can accept a variety of substrates, including small-molecule fluorescent dyes that significantly outperform fluorescent proteins for live imaging^44–46^. We included a Hygromycin resistance marker in our HaloTag constructs, enabling their use in cells that already carry the Puromycin resistance marker that was used to select mNG knock-ins. Our mNG and HaloTag knock-in donors utilize the same frame selector Cas9/sgRNA plasmids, so that a tagging strategy that has been demonstrated for one tag can be re-used to insert a different tag.

**Figure 5:**
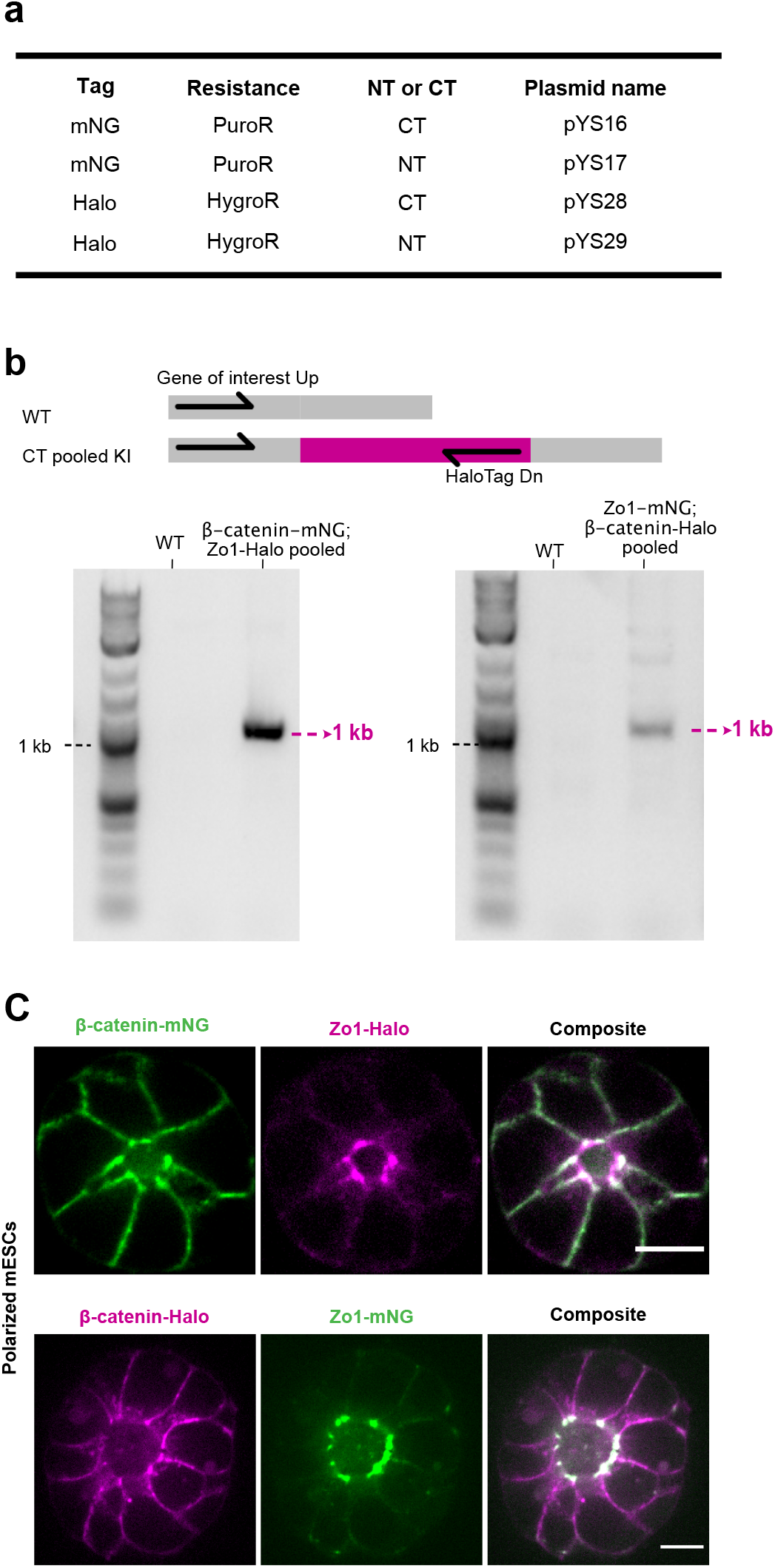
Dual-color knock-in by sequential transfection. **5a.**Table summarizing donor vectors for either mNG or HaloTag insertion at either terminus of a protein. **5b**. PCR genotyping of β-catenin-mNG; Zo1-Halo and Zo1-mNG; β-catenin-Halo pooled cells. **5c**. Multicolor imaging of dual-tagged cell lines after epithelial polarization in 3D culture. Scale bars represent 10 µm.

To demonstrate the utility of our Halo tagging vectors for two-color knock-in, we constructed dual-labeled cell lines carrying tags on both β-catenin and Zo1. We generated Zo1-HaloTag in a β-catenin-mNG background, and β-catenin-HaloTag in a Zo1-mNG background. After drug selection, PCR genotyping confirmed a correct insertion in both cell lines (Figure 5b). We then incubated cells with the JFX_646_ far-red fluorescent HaloTag ligand^47^ and performed live imaging in 3D culture. Both dual-labeled cell lines showed the expected distributions of β-catenin and Zo1 at the adherens and tight junctions, respectively (Figure 5c). In summary, our method can label more than one protein with different FPs in the same cell line by sequential transfection, which makes it possible to evaluate co-localization of endogenous proteins in living cells. In the future, it should be possible to achieve multi-color labeling by adding additional FPs and selectable markers to our existing design.

## Discussion

We have demonstrated an efficient and rapid protocol for FP knock-in in mammalian stem cells. Five distinct loci have been edited with successful insertion on the first attempt, highlighting the generality and ease of the strategy. Our method requires minimal cloning since only the genespecific sgRNA vector needs to be customized, and it offers flexibility to tag either end of a gene to produce functional proteins for live imaging.

Our workflow for FP knock-in in mESCs is simple and efficient (Figure 6). It takes less than 1 hour to design an sgRNA vector. Within one week, the sgRNA vector can be cloned and minicircle donors can be produced from the premade parent donor plasmids. Transfection into mESCs takes less than 1 hour. On the second day after transfection, drug treatment is used to enrich in-frame edited cells for 2-3 days. After drug selection, mESCs can be expanded for imaging, PCR genotyping, and maintenance within 1-2 weeks. After validation by PCR genotyping, monoclonal cell lines can be picked by seeding low cell numbers in a 10cm plate for 1 week. Then the monoclonal cell lines can be expanded for PCR genotyping and sequencing. Depending on cell proliferation rate, this can take 2-3 weeks; thus, the total time required to isolate monoclonal knock-in cell lines is still reasonable at 5-6 weeks. In the supplemental material, we provide a detailed protocol for streamlined FP knock-in.

**Figure 6:**
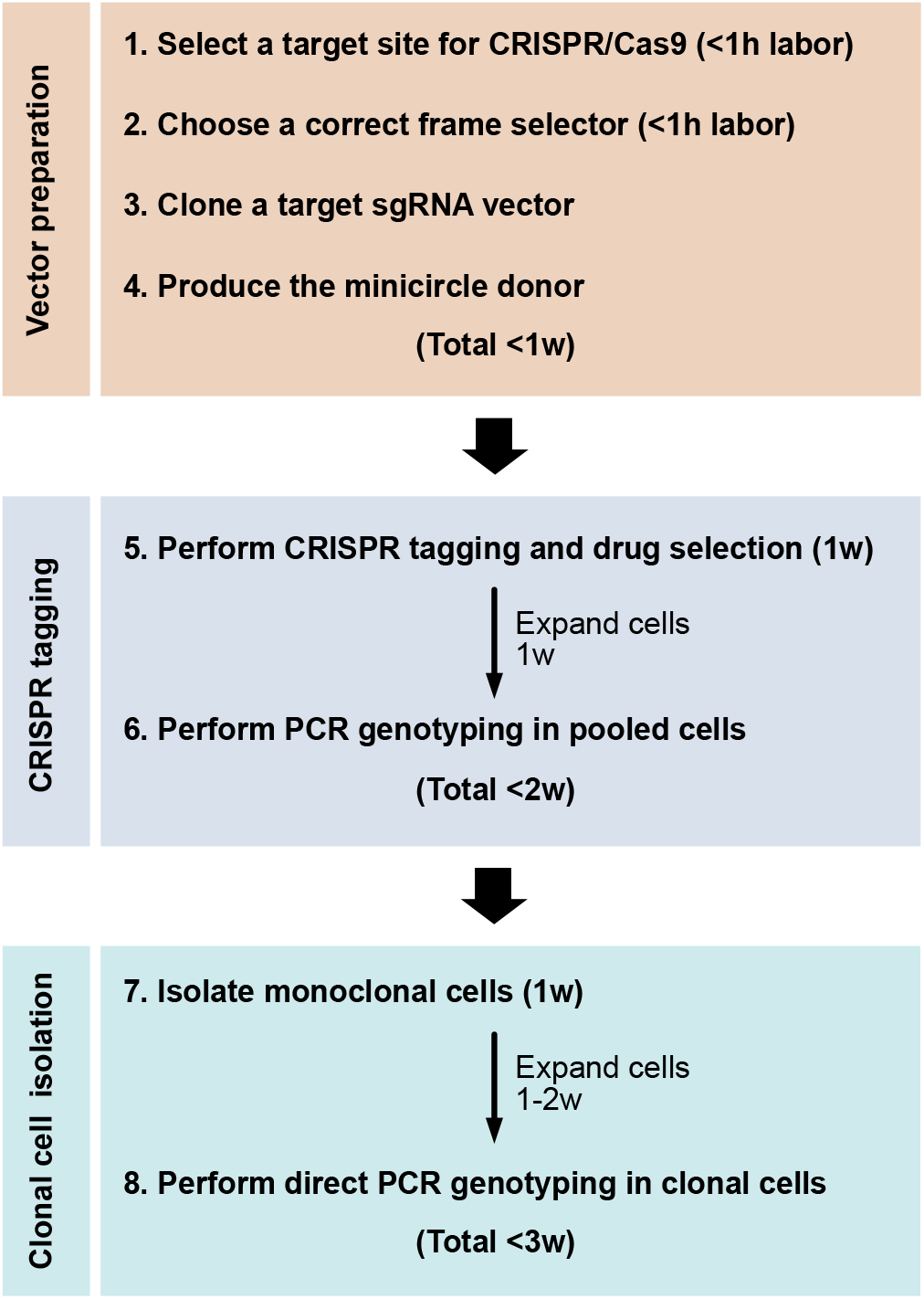
CRISPR tagging workflow.

Combining NHEJ and drug selection gives rise to a high proportion (>75%) of functional knock-ins. The 2-week timetable from transfection to imaging is comparable to the time required for establishing a stable cell line expressing an exogenous transgene, but endogenous tagging can avoid overexpression artifacts and thus provides more reliable results. The high insertion efficiency facilitates isolation of monoclonal cell lines when needed, since fewer than 12 clones are sufficient to obtain multiple functional knock-in clones. It is feasible to isolate seamless and bi-allelic insertions, though this may require screening more clones.

A potential future application of this approach is to enable rapid generation of knock-in mice expressing fluorescent proteins from endogenous loci. While current approaches for gene tagging in mice via zygote or 2-cell embryo injection can be highly efficient^48,49^, this varies based on the user as well as the locus being targeted. In addition, one must then characterize the generated insertions in mice, which are often mosaic due to perdurance of the CRISPR reagents past the stage at which they are injected^50–52^. By rapidly editing mESCs with positive selection, characterizing them in culture and then injecting them into blastocysts to generate chimeras, it should be feasible to establish new mouse lines or to perform microscopy studies in the chimeric mice themselves.

It is important to note that, although we observed indels or mutations at the junctions between these targeted genes and the donor, small indels do not significantly undermine the utility of this approach. Since the donor vector has no constitutive promoter, after insertion, the FP and drug resistance marker can only be expressed by the target gene’s native promoter. Drug selection thus ensures that surviving cells have a correct knock-in, with the insertion in-frame and in proper orientation. Indels and seamless insertions most likely result from two distinct NHEJ pathways, aNHEJ and cNHEJ, respectively. aNHEJ is end resection-dependent and thus produces small deletions, while cNHEJ is end resection-independent and can produce seamless insertions^11,34^. In the future, it may be possible to increase the frequency of seamless insertions by inhibiting end resection / aNHEJ. Alternatively, by careful screening, it is feasible to achieve scarless insertion if necessary.

Compared to an overexpression strategy, endogenous gene tagging is expected to recapitulate the native expression pattern of the protein of interest more faithfully. However, a caveat is that endogenous tagging still might compromise the function of some proteins, either by influencing gene expression or by steric effects on protein function. For example, our mono-allelic mNG-Myh9 clones had indels at the 5’ end of the untagged allele, some of which may disrupt the expression of the untagged allele. For haploinsufficient genes, such an effect could cause unwanted phenotypes. Additionally, the tagged alleles of β-catenin-mNG that we generated appear to be expressed at lower levels than the untagged protein, for reasons that are unclear. Although this did not prevent us from observing the localization of β-catenin-mNG in microscopy experiments, it could present an issue for experiments where a quantitatively normal level of tagged protein is critical. Thus, the utility of cells carrying endogenously tagged proteins needs to be assessed on a case-by-case basis and will depend on the goal of the experiment.

Our strategy is expected to be less efficient when tagging silent, non-expressed genes compared to the actively transcribed genes we have tested here. Some genes relevant to stem cell biology and differentiation, such as Otx2, Sox11 and Oct6, are not expressed in naïve mESCs. Because the FP and drug resistance sequences rely on the native promoter for expression in our system, the drug resistance marker will not be expressed if the target gene is not expressed. However, NHEJ has been proven efficient at repairing DSBs in both active and silent genes^18^. Thus, a possible workaround for a silent gene would be to target the gene in naïve pluripotent stem cells, pick clones, and then generate duplicates of each clone in 96-well plates. One copy would be screened for insertions, either by PCR or by differentiating cells and performing drug selection, and then positive clones could be retrieved from the master plate.

Alternatively, it might be possible to select for insertions under culture conditions that induce upregulation of some genes that are silent in naïve mESCs, but that are still permissive for reversion to the naïve state upon culturing in 2i/LIF^53–55^. A second limitation of our method is the inability of this approach to tag a gene internally, rather than at the N- or C-terminus. The CRISPR-mediated insertion of exon (CRISPIE) approach is probably a better choice for internal tags^24^. The CRISPIE system^24^ inserts an FP coding sequence flanked by splicing signals into an intronic location of the desired gene so that the insertion junctions will be spliced out, leaving the protein coding sequence unaffected by indels caused by NHEJ. This approach is well-suited for internal FP insertion but requires extra cloning to produce N- or C-terminal tags.

Despite these limitations, our protocol holds promise for a fast, easy, robust endogenous tagging in mammalian cells, allowing native protein visualization in living cells without overexpression.

## Methods

### Plasmid information

A list of plasmids generated for this study is provided in Table S1. All plasmids will be deposited at Addgene upon manuscript acceptance.

### Target Cas9/sgRNA vector cloning

pDD428, which contains the U6 promoter to express *Myh9* C-terminal sgRNA and the EF1α promoter to express mCherry-Cas9, was used as a template for new target sgRNA vector construction. We tested two alternative cloning strategies (see Supplemental Protocol for details). First, we used the NEB site-directed mutagenesis kit, following the manufacturer’s instructions. The entire pDD428 vector was PCR amplified with a forward primer that included the new target protospacer in place of the existing protospacer, and a reverse primer that annealed to the U6 promoter. The PCR product was self-ligated, and clones were verified by Sanger sequencing. In the second approach, we generated a short PCR fragment by amplifying pDD428 with a forward primer that included the new target protospacer and a universal reverse primer that annealed to the EF1α promoter. The vector backbone was prepared by digesting pDD428 with SalI-HF and EcoRI-HF, and NEB HiFi DNA assembly was performed to insert the PCR product containing the new protospacer into the digested vector. Table S2 provides primers used for each strategy, and Table S3 lists the target protospacers used in our sgRNA vectors.

### Minicircle vector production

We have generated four parent donor vectors: mNG-PuroR and Halo-HygroR for C-terminal tagging, and PuroR-mNG and HygroR-Halo for N-terminal tagging (Figure 5a). These constructs are based on an empty parent minicircle cloning vector (MN100A-1; System Biosciences (SBI), Palo Alto, CA). The premade parent donor vectors contain attB-attP recombination sites flanking the FP plus drug-resistant gene cassette. They have been transformed into an engineered *E. coli* called ZYCY10P3S2T (SBI MN900A-1), which enables minicircle donor production^36^. To make minicircle donors, fresh ZYCY10P3S2T *E. coli* carrying a parent donor vector were grown in 50 mL of TB medium containing 50 µg/mL Kanamycin at 30°C with shaking at 250 rpm until the OD value was 4-6. When the proper OD value was achieved, 50 mL of induction buffer was added and growth was continued at 30°C for 3h to produce minicircles, then at 37°C for 1h to degrade the vector backbone. The induction buffer was made by mixing 400 mL of fresh LB, 16 mL of 1N NaOH and 0.4 mL 20% L-arabinose (Thermo Fisher AC104980250). Finally, we collected and aliquoted 100 mL induced *E. coli* into 3 mL per tube. The induced *E. coli* was spun down at 4°C. After removing media, we stored the pellets of induced *E. coli* at -20°C. To extract minicircle donors, we used the QIAprep Spin Miniprep Kit (Qiagen 27104). Before minicircle donor extraction, we thawed one aliquot of induced *E. coli* at 4°C and resuspended the pellet with 500 μL of P1 buffer. Then we lysed it with 500 μL of P2 buffer for 5 min and mixed it with 700 μL of N3 buffer, centrifuged and transferred supernatant to one QIAprep spin column. We washed the spin column with 500 μL of PB buffer and 750 μL of PE buffer. Finally, we eluted minicircle donors with 50 μL of Nuclease-free water. 3 mL of induced *E. coli* yielded around 2.5 µg of minicircle donors (50 ng/μL in 50 μL), which is sufficient for 6 transfections. The quality of the minicircle donor was checked by linearizing the donor and performing gel electrophoresis (See Figure S2b). See Supplemental Protocol for details.

### mESC culture

J1 mESCs (ATCC, SCRC-1010) were maintained in 2i/LIF medium at 37 °C, 7% CO_2_ in gelatin-coated plates^**29**^. 2i/LIF was made of N2B27 with 3 µM of GSK3β inhibitor (Sigma SML1046), 1 µM of MEK inhibitor (Sigma PZ0162) and 100 U/mL of Leukemia Inhibitory Factor (Sigma ESG1106). N2B27 was either purchased as a commercial product (ESGRO Complete Basal Medium, Sigma SF002-500) or made in a 1:1 mixture by 487 mL of DMEM/F12 (Sigma D6421) and 487 mL of Neurobasal (Thermo Fisher 21103049) supplemented with 0.5X B27(Invitrogen 17504044), 1X N2 (homemade, see below), 50 μM β-mercaptoethanol (Sigma M3148-25ML), and 2 mM L-glutamine (Thermo Fisher 25030081). Homemade N2 contained DMEM/F12 (Sigma D6421), 2.5 mg/mL insulin (Sigma I9278), 10 mg/mL Apo-transferrin (Sigma T1147), 0.75% bovine albumin fraction V (Thermo Fisher 15260037), 1.98 μg/mL progesterone (Sigma P8783-1G), 1.6 mg/mL putrescine dihydrochloride (Sigma P5780-5G) and 0.518 μg/mL sodium selenite (Sigma S5261-10G). The final density for cell seeding was 15 × 10^4 cells/mL. 2i/LIF was renewed every day and mESCs were routinely passaged every 2-3 days. Mycoplasma testing (Southern Biotech 13100-01) was performed periodically.

For routine passaging of mESCs in gelatin-coated 10cm plate, the growth medium 2i/LIF was removed and 3.5 mL Accutase (Sigma SCR005) was added to cells and incubated at 37 °C for 5 min. To dissociate mESCs into single cell suspension, cells were pipetted up and down 10-15 times. Then, the cell suspension was added to 10.5 mL of wash medium (500 mL of DMEM/F12 + 8 mL of bovine albumin fraction V) in a 15 mL tube. Cells were spun down for 5 min at 1000 rpm, and 1 mL of 2i/LIF was used to resuspend cells. 10 µL cells were mixed with 10 μL of Trypan blue (Thermo Fisher 15250061) and counted using a cell counter (Invitrogen AMQAF1000). Finally, 150 × 10^6 cells with 10 mL of 2i/LIF were seeded in a new gelatin-coated 10 cm plate.

### mESC 3D culture

Growth factor reduced Matrigel (Corning 356230) was used in 3D spheroid cultures^40,41^. One well of a 15-well imaging plate (ibidi 81506) was covered with 1 µL of ice-cold Matrigel and incubated for 15 min at 37 °C to allow the solidification of Matrigel. N2B27 or 2i/LIF was used to resuspend mESCs and cell density was adjusted to 10 × 10^4 cell/mL. Then 30 µL of cells (0.3 × 10^4 cell per well) were seeded on each Matrigel-coated well. When the cells had settled down to the Matrigel (15 min after seeding), the media were removed and replaced with 50 µL of N2B27+M or 2i/LIF+M (N2B27 or 2i/LIF containing 5% Matrigel). It took 60-72h for cells in N2B27+M to form polarized spheres. Cells in 2i/LIF remained unpolarized throughout 3D culture.

### Co-transfection

We transfected three plasmids (an FP donor vector, a frame selector Cas9/sgRNA vector and a target Cas9/sgRNA vector) into mESCs using Lipofectamine 2000 (Thermo Fisher 11668027). The molar ratio of [donor vector] : [frame selector] : [target sgRNA] plasmid was typically 20:1:1, and was achieved using 364 ng minicircle donor,130 ng target Cas9/sgRNA vector, and 130 ng frame selector. The three plasmids were mixed and diluted to 50 µL in Opti-MEM Medium (Gibco 31985062). 1.25 µL Lipofectamine 2000 was diluted to 50 µL in Opti-MEM Medium, then added to the 50 μL of DNA solution and incubated at room temperature for 5 min. After incubation, the 100 μL DNA/Lipofectamine 2000 mixture was transferred to a gelatin-coated 12-well plate. 5 × 10^5 single cells in suspension per well were added to DNA/Lipofectamine 2000 mixture. 2i/LIF was added up to 1 mL per well. The same procedures were performed with 50 μL of pure Opti-MEM Medium and 50 μL of diluted Lipofectamine 2000 as a negative control. We replaced the transfection mixture with 2-3 mL of fresh 2i/LIF and observed mCherry-Cas9 signal in the transfected cells on the next day. Triplicates of co-transfection were used for each gene tagging. Table S4 lists plasmids that were co-transfected in each experiment.

### Drug selection

On day 2 after transfection, we treated the transfected cells with either 1-2 µg/mL Puromycin or 100-200 µg/mL Hygromycin, depending on the chosen donor vector. Each day, the plates were gently washed 1-2 times with wash medium to remove dead cells, and then fresh drug-containing medium was added. This process was repeated daily for 2-3 days of drug treatment. Most of the cells appeared floating and dead after 1 day of Puromycin or 3 days of Hygromycin treatment. After selection was complete, the medium was replaced with 2i/LIF to expand surviving cells. On day 6 or 7 after transfection, the drug-resistance cells started to form small colonies. The surviving cells were maintained in 2i/LIF and were ready for genomic DNA extraction or imaging within 14 days after transfection.

### Isolation of monoclonal cells

To generate monoclonal cell lines, 1 × 10^4 pooled edited cells with 10 mL of 2i/LIF were seeded on gelatin-coated10 cm plates. 2i/LIF was renewed every day until the individual cell formed a discernible colony that was visible with a naked eye. On d7-d10, colonies were picked using a P10 pipet set at 5 µL under a dissection microscope. We incubated the picked colonies in a U-shape-bottom 96-well plates with 20 µL Accutase in each well at 37°C for 10 min. After incubation, we added 150 µL of 2i/LIF to the Accutase/cell mixture, pipetted the mixture up and down to dissociate colonies into single cells, then transferred cell suspension to a gelatin-coated U-shape-bottom 96-well plate. On the next day, we gently removed most of the medium but left enough to cover the cells and added fresh 150 µL of 2i/LIF. We changed 2i/LIF every day until cells reached confluence. If cells didn’t attach to the bottom of the plate, 1% FBS in 2i/LIF was added to help cell adhesion.

### Genomic DNA extraction

After transfecting and enriching drug-resistant cells, we split putatively edited cells for genomic DNA extraction. To extract genomic DNA from mESCs, we used the PureLink Genomic DNA Mini Kit (Thermo Fisher K182001 by following the manufacturer’s instructions. Typically, 5 × 10^5 mESCs yielded around 6.25 µg of genomic DNA (approximately 250 ng/μL in 25 μL). To verify insertion in pooled cells, we extracted genomic DNA of pooled cells for PCR genotyping. Meanwhile, we collected genomic DNA of WT cells as a negative control.

### Quick lysis

For monoclonal cells, genomic DNA Mini Kit can be used for PCR genotyping. However, it was expensive and time-consuming to genotype many clonal cell lines in this way. We therefore used quick lysis in place of genomic DNA extraction for clonal cells. When clonal cells reached confluence in the 96-well plate, cells were split into two sets of plates, one in a 96-well plate for PCR genotyping, the other in a 24-well plate for subculture. We changed 2i/LIF every day until cells in the 96-well plate reached confluence. Then we gently removed all the medium in the 96-well plate and added 100 µL of Quick lysis buffer. Quick lysis buffer comprised 10 mg/mL proteinase K powder (Sigma P6556), 50 mM KCl, 10 mM Tris-HCl pH 8, 2 mM MgCl_2_, 1% (v/v) NP40 (Thermo Fisher 85124), 0.45%(v/v) Tween 20 and milli-Q water. We pipetted up and down to dissociate colonies, then transferred the cell lysates to PCR Tube Strips, sealed the plate and incubated at 65°C for 2h followed by 95°C for 10 min. Finally, we diluted cell lysates 1:20 with nuclease-free water and used 1 µL of the resulting solution for PCR genotyping.

### PCR genotyping

To verify insertions, we used a forward primer upstream of the cutting site and a reverse primer inside the donor vector. For C-terminal insertions, we used a reverse primer inside the mNG or HaloTag coding sequence. For N-terminal insertion, the reverse primer was inside the PuroR coding sequence. To distinguish whether an insertion is mono-allelic or bi-allelic (i.e., only one genomic copy or both genomic copies are tagged), we used a pair of primers upstream and downstream of the cutting site. Table S5 lists primer information for PCR genotyping. (See Supplemental Protocol for details)

As template for PCR genotyping, we used 50 ng of purified genomic DNA from pooled edited or WT cells or 1 µL diluted cell lysate from clonal cells. PCR was performed using LongAmp Taq 2X Master Mix (NEB M0287S) according to the manufacturer’s instructions, and results were visualized by electrophoresis.

### Sanger sequencing of insertions

After verification by PCR and electrophoresis, we extracted specific bands using the Zymoclean Gel DNA Recovery Kits (Zymo D4007/D4008). Then we submitted purified amplicons with a corresponding forward primer or reverse primer for Sanger sequencing (Table S5). We aligned Sanger sequencing results with WT or knock-in allele to validate insertion and characterize the insertion sequence.

### Western blotting

Cells were resuspended from 2D culture with PBS at various cell concentrations. After lysing with NuPAGE LDS Sample Buffer (Thermo-Fisher NP0007) containing 100 mM DTT, the reduced cell samples were incubated at 70°C for 10 min. Equal amounts of cell sample and 1x benzonase nuclease (Sigma 70746-3) was added to degrade nucleic acids. Samples were sonicated to further reduce any contamination of nucleic acids. The samples were run on a 4-12% Bis-Tris NuPAGE gel (Thermo-Fisher NP0323) using MES Running buffer (Thermo-Fisher NP0002) at 160V for 1h with antioxidant. The gel was then transferred to a polyvinylidene fluoride (PVDF) membrane at 20V for 1h. A blocking buffer (5% non-fat dry milk dissolved in Phosphate Buffered Saline-Tween (PBST) was used for 0.5h at room temperature to prevent non-specific binding. The membrane was then incubated with primary antibodies overnight at 4°C: anti-Alpha Tubulin (1:200, DSHB 12G10) and *β*-catenin (1:10,000 dilution, Sigma ABE208). The membrane was washed three times using PBST, blocked using 5% non-fat dry milk in PBST, and then incubated with secondary antibodies for 45 min at room temperature: anti-Mouse 680 (1:10,000 dilution, Thermo-Fisher A-21057) and anti-Rabbit 800 (1:20,000 dilution, Thermo-Fisher SA5-10036). The membrane was washed three times with PBST followed by two washes with PBS. The membrane was imaged using a LICOR-Odyssey CLx.

### Cell dye incubation

To image cells with HaloTag, HaloTag ligand dye JFX_646_ (gift from Luke Lavis) was used. It was firstly dissolved in acetonitrile to 1 mM and aliquoted into 2 µL in PCR tubes. The dye was dried with a vacuum centrifuge for storage at -20°C in a desiccator. Before imaging, we dissolved one aliquot with 2 µl of DMSO to reconstitute 1mM dye. Then we diluted it into 0.037 µM with growth medium. Before imaging, we incubated cells with 0.037 µM JFX_646_+2i/LIF for at least 30 min in a 15-well imaging plate at 37°C.

Cell tracing dyes including SPY555-Actin (Cytoskeleton CY-SC202), SPY650-Tubulin (Cytoskeleton CY-SC503) and SPY650-DNA (Cytoskeleton CY-SC501) were dissolved in 50 µL of fresh DMSO to reconstitute 1000X. Then they were aliquoted into 5 µl in PCR tubes, protected from light and stored at -20°C. Before imaging, we incubated cells with 1X cell tracing dye with growth medium for at least 15 min in a 15-well imaging plate at 37°C. Then cells were ready for imaging. There was no wash step after dye incubation because all dyes were fluorogenic and produced low background fluorescence.

### Confocal live microscopy

Cells were seeded in 15-cell imaging slides and kept in the tissue culture incubator until ready for imaging. To observe epithelial polarity in 3D culture, cells were incubated for 60-72h. To observe the frequency of GFP+ cells in 2D culture, cells were incubated for 12h. Before imaging, a humidifier was filled with water, and a gas mixer (Okolab 2GF-MIXER) with air pump and 100% CO_2_ and temperature controller (Okolab H401-T-CONTROLLER) were turned on so that the imaging chamber (Okolab H201-NIKON-TI-S-ER) could pre-equilibrate to the correct humidity, 7% CO_2_, and 37°C for live cells. Then images were acquired using one of two microscopes. The images shown in Figure 1 were acquired using a Nikon Eclipse Ti-2 microscope equipped 20X, 0.75 NA and 60X, 1.45 NA objectives; a Photometrics PrimeBSI camera; an OptoSpin filter wheel (CAIRN Research, Kent, England), and an vt-iSIM super-resolution confocal scan head (VisiTech international, Sunderland, UK). mNeonGreen fluorescence was excited using a 505 nm diode laser. All other images were acquired using a Nikon Eclipse Ti-2 microscope equipped 20X, 0.75 NA and 60X, 1.4 NA objectives; an 89 North LDI-7 laser diode illuminator; a Photometrics Prime95B camera; and a Crest X-Light V3 spinning disk confocal head. mNG, SPY555-Actin, SPY650-Tubulin SPY650-DNA and HaloTag-JF_646_ were excited using the appropriate lines of the LDI-7 illuminator. For fixed imaging, mESCs were imaged by multiple z-planes with a step of 1.5 µm. For Timelapse live imaging, cells were imaged every 5-30 min in multiple z-planes with a step of 1.5 µm.

To prepare figures, the midplane of confocal image stacks was selected, and images were cropped and rotated using FIJI. Some images were processed with a Gaussian Blur filter with a 0.7 pixel radius to reduce background noise, and the brightness and contrast were adjusted for visibility of the signals. No other image manipulations were performed.

### Cell cryopreservation and recovery

After genotyping, we froze pooled and at least 5 different monoclonal cells. To freeze edited cells for future use, cell number was counted and at least 5 × 10^6 cells were collected in 2i/LIF per tube. Then cells were spun down for 5 min at 1000 rpm and resuspended cells by using 500 μL of freezing medium containing 450 μL of 2i/LIF and 50 μL of DMSO per tube. Cells were transferred in a pre-labeled cryovial and put the cryovial into a freezing chamber and placed in a -80°C freezer immediately. Finally, the cryovial was transferred from the freezing chamber to the -80°C freezer or a liquid nitrogen tank.

To recover edited cells in cryovial for imaging or sequential transfection, the frozen cells were thawed in 37°C water bath for 1.5 min, then mixed with 10 mL wash medium and transferred into a 15 mL tube. Cells were spun down for 5 min at 1000 rpm, resuspended with 10 mL of 2i/LIF and seeded into a 10cm gelatin-coated plate. Cells were passaged at least three times before any experiments.

## Supporting information

Supplemental Protocol

## Acknowledgements

This work is supported by the U.S. National Institutes of Health (R01 GM138443) and a CPRIT Scholar award from the Cancer Prevention and Research Institute of Texas (RR170054) to DJD. We thank Ivy Chang and Qiuxia Zhao for help with cloning; Jonghwan Kim, Steve Vokes and members of the Dickinson lab for helpful discussions and comments on the manuscript; Luke Lavis for sharing JaneliaFluor dyes; and Xiangjun Zhao (University of Manchester) for sharing the quick lysis protocol.

## Author Contributions

Y.S. and D.J.D. conceived the project and designed the experiments. Y.S., N.K., L.O. and D.J.D. constructed plasmids and prepared minicircle donors. N.K. and L.O. assisted with cell culture and genotyping, and N.K. performed western blotting experiments. Y.S. performed all other experiments, analyzed the data, and prepared the figures. Y.S. and D.J.D. wrote the manuscript text, and all authors discussed and contributed to the final version. D.J.D. supervised the project and secured funding.

## References

1. Gibson, T. J., Seiler, M. & Veitia, R. A. The transience of transient overexpression. Nat Methods 10, 715–721 (2013).

2. Wach, A., Brachat, A., Alberti-Segui, C., Rebischung, C. & Philippsen, P. Heterologous HIS3 Marker and GFP Reporter Modules for PCR-Targeting in Saccharomyces cerevisiae. Yeast 13, 1065–1075 (1997).

3. Huh, W.-K. et al. Global analysis of protein localization in budding yeast. Nature 425, 686–691 (2003).

4. Dickinson, D. J., Ward, J. D., Reiner, D. J. & Goldstein, B. Engineering the Caenorhabditis elegans genome using Cas9-triggered homologous recombination. Nature Methods 10, 1028–1034 (2013).

5. Dickinson, D. J., Pani, A. M., Heppert, J. K., Higgins, C. D. & Goldstein, B. Streamlined Genome Engineering with a Self-Excising Drug Selection Cassette. Genetics 200, 1035–1049 (2015).

6. Paix, A., Folkmann, A., Rasoloson, D. & Seydoux, G. High Efficiency, Homology-Directed Genome Editing in Caenorhabditis elegans Using CRISPR-Cas9 Ribonucleoprotein Complexes. Genetics 201, 47–54 (2015).

7. Dokshin, G. A., Ghanta, K. S., Piscopo, K. M. & Mello, C. C. Robust Genome Editing With Short Single-Stranded and Long, Partially Single-Stranded DNA Donors in Caenorhabditiselegans. Genetics (2018).

8. Baena-López, L. A., Alexandre, C., Mitchell, A., Pasakarnis, L. & Vincent, J.-P. Accelerated homologous recombination and subsequent genome modification in Drosophila. Development 140, 4818–4825 (2013).

9. Li-Kroeger, D. et al. An expanded toolkit for gene tagging based on MiMIC and scarless CRISPR tagging in Drosophila. eLife 7, e1002472 (2018).

10. Bukhari, H. & Müller, T. Endogenous Fluorescence Tagging by CRISPR. Trends Cell Biol 29, 912–928 (2019).

11. Chiruvella, K. K., Liang, Z. & Wilson, T. E. Repair of Double-Strand Breaks by End Joining. Csh Perspect Biol 5, a012757 (2013).

12. Ran, F. A. et al. Genome engineering using the CRISPR-Cas9 system. Nat Protoc 8, 2281–2308 (2013).

13. Yang, L., Yang, J. L., Byrne, S., Pan, J. & Church, G. M. CRISPR/Cas9-Directed Genome Editing of Cultured Cells. Curr Protoc Mol Biology 107, 31.1.1-31.1.17 (2014).

14. Haupt, A. et al. Endogenous Protein Tagging in Human Induced Pluripotent Stem Cells Using CRISPR/Cas9. J Vis Exp Jove 58130 (2018) doi:10.3791/58130.

15. Morrow, C. S., Porter, T. J. & Moore, D. L. Fluorescent tagging of endogenous proteins with CRISPR/Cas9 in primary mouse neural stem cells. Star Protoc 2, 100744 (2021).

16. Mao, Z., Bozzella, M., Seluanov, A. & Gorbunova, V. DNA repair by nonhomologous end joining and homologous recombination during cell cycle in human cells. Cell Cycle 7, 2902–2906 (2008).

17. Mao, Z., Bozzella, M., Seluanov, A. & Gorbunova, V. Comparison of nonhomologous end joining and homologous recombination in human cells. Dna Repair 7, 1765–1771 (2008).

18. He, X. et al. Knock-in of large reporter genes in human cells via CRISPR/Cas9-induced homology-dependent and independent DNA repair. Nucleic Acids Res 44, e85–e85 (2016).

19. Bachu, R., Bergareche, I. & Chasin, L. A. CRISPR-Cas targeted plasmid integration into mammalian cells via non-homologous end joining. Biotechnol Bioeng 112, 2154–2162 (2015).

20. Suzuki, K. et al. In vivo genome editing via CRISPR/Cas9 mediated homology-independent targeted integration. Nature 540, 144–149 (2016).

21. Lackner, D. H. et al. A generic strategy for CRISPR-Cas9-mediated gene tagging. Nat Commun 6, 10237 (2015).

22. Schmid-Burgk, J. L., Höning, K., Ebert, T. S. & Hornung, V. CRISPaint allows modular base-specific gene tagging using a ligase-4-dependent mechanism. Nat Commun 7, 12338 (2016).

23. Artegiani, B. et al. Fast and efficient generation of knock-in human organoids using homology-independent CRISPR–Cas9 precision genome editing. Nat Cell Biol 22, 321–331 (2020).

24. Zhong, H. et al. High-fidelity, efficient, and reversible labeling of endogenous proteins using CRISPR-based designer exon insertion. Elife 10, e64911 (2021).

25. Zeng, F. et al. A Simple and Efficient CRISPR Technique for Protein Tagging. Cells 9, 2618 (2020).

26. Zhong, H. et al. High-fidelity, efficient, and reversible labeling of endogenous proteins using CRISPR-based designer exon insertion. Elife 10, e64911 (2021).

27. Atkins, J. F. et al. A case for “StopGo”: Reprogramming translation to augment codon meaning of GGN by promoting unconventional termination (Stop) after addition of glycine and then allowing continued translation (Go). Rna 13, 803–810 (2007).

28. Doronina, V. A. et al. Site-Specific Release of Nascent Chains from Ribosomes at a Sense Codon. Mol Cell Biol 28, 4227–4239 (2008).

29. Mulas, C. et al. Correction: Defined conditions for propagation and manipulation of mouse embryonic stem cells (doi:10.1242/dev.173146). Development 146, dev178970 (2019).

30. Czechanski, A. et al. Derivation and characterization of mouse embryonic stem cells from permissive and nonpermissive strains. Nat Protoc 9, 559–574 (2014).

31. Chung, S. et al. Analysis of Different Promoter Systems for Efficient Transgene Expression in Mouse Embryonic Stem Cell Lines. Stem Cells 20, 139–145 (2002).

32. Dean, S. O., Rogers, S. L., Stuurman, N., Vale, R. D. & Spudich, J. A. Distinct pathways control recruitment and maintenance of myosin II at the cleavage furrow during cytokinesis. Proc National Acad Sci 102, 13473–13478 (2005).

33. Labun, K. et al. CHOPCHOP v3: expanding the CRISPR web toolbox beyond genome editing. Nucleic Acids Res 47, W171–W174 (2019).

34. Ceccaldi, R., Rondinelli, B. & D’Andrea, A. D. Repair Pathway Choices and Consequences at the Double-Strand Break. Trends Cell Biol 26, 52–64 (2016).

35. Deriano, L. & Roth, D. B. Modernizing the Nonhomologous End-Joining Repertoire: Alternative and Classical NHEJ Share the Stage. Annu Rev Genet 47, 433–455 (2013).

36. Kay, M. A., He, C.-Y. & Chen, Z.-Y. A robust system for production of minicircle DNA vectors. Nat Biotechnol 28, 1287–1289 (2010).

37. Snapp, E. Design and Use of Fluorescent Fusion Proteins in Cell Biology. Curr Protoc Cell Biology 27, 21.4.1-21.4.13 (2005).

38. Tikhonov, M. V., Maksimenko, O. G., Georgiev, P. G. & Korobko, I. V. Optimal artificial mini-introns for transgenic expression in the cells of mice and hamsters. Mol Biol+ 51, 592–595 (2017).

39. Rodriguez, J. M. et al. APPRIS: annotation of principal and alternative splice isoforms. Nucleic Acids Res 41, D110–D117 (2013).

40. Bedzhov, I. & Zernicka-Goetz, M. Self-organizing properties of mouse pluripotent cells initiate morphogenesis upon implantation. Cell 156, 1032–1044 (2014).

41. Shahbazi, M. N. et al. Pluripotent state transitions coordinate morphogenesis in mouse and human embryos. Nature 122, 881 (2017).

42. Bryant, D. M. & Mostov, K. E. From cells to organs: building polarized tissue. Nat Rev Mol Cell Biol 9, 887–901 (2008).

43. Choi, H.-J., Huber, A. H. & Weis, W. I. Thermodynamics of beta-catenin-ligand interactions: the roles of the N- and C-terminal tails in modulating binding affinity. 281, (2006).

44. Los, G. V. et al. HaloTag: a novel protein labeling technology for cell imaging and protein analysis. ACS chemical biology 3, 373–382 (2008).

45. Grimm, J. B. et al. A general method to improve fluorophores for live-cell and single-molecule microscopy. Nat Methods 12, 244–250 (2015).

46. Grimm, J. B. et al. A general method to optimize and functionalize red-shifted rhodamine dyes. Nat Methods 17, 815–821 (2020).

47. Grimm, J. B. et al. A General Method to Improve Fluorophores Using Deuterated Auxochromes. Jacs Au 1, 690–696 (2021).

48. Ge, X. A. & Hunter, C. P. Efficient Homologous Recombination in Mice Using Long Single Stranded DNA and CRISPR Cas9 Nickase. G3 Genes Genomes Genetics 9, 281–286 (2018).

49. Gu, B., Posfai, E. & Rossant, J. Efficient generation of targeted large insertions by microinjection into two-cell-stage mouse embryos. Nature Biotechnology 36, 632–637 (2018).

50. Yen, S.-T. et al. Somatic mosaicism and allele complexity induced by CRISPR/Cas9 RNA injections in mouse zygotes. Dev Biol 393, 3–9 (2014).

51. Oliver, D., Yuan, S., McSwiggin, H. & Yan, W. Pervasive Genotypic Mosaicism in Founder Mice Derived from Genome Editing through Pronuclear Injection. Plos One 10, e0129457 (2015).

52. Mehravar, M., Shirazi, A., Nazari, M. & Banan, M. Mosaicism in CRISPR/Cas9-mediated Genome editing. Dev Biol 445, 156–162 (2018).

53. D’Aniello, C. et al. Vitamin C and l-Proline Antagonistic Effects Capture Alternative States in the Pluripotency Continuum. Stem Cell Rep 8, 1–10 (2017).

54. Neagu, A. et al. In vitro capture and characterization of embryonic rosette-stage pluripotency between naive and primed states. Nat Cell Biol 22, 534–545 (2020).

55. Kinoshita, M. et al. Capture of Mouse and Human Stem Cells with Features of Formative Pluripotency. Cell Stem Cell 28, 453-471.e8 (2021).

