## Supplemental Figures for "Efficient and rapid fluorescent protein knock-in with universal donors in mammalian stem cells"

### Figure S1

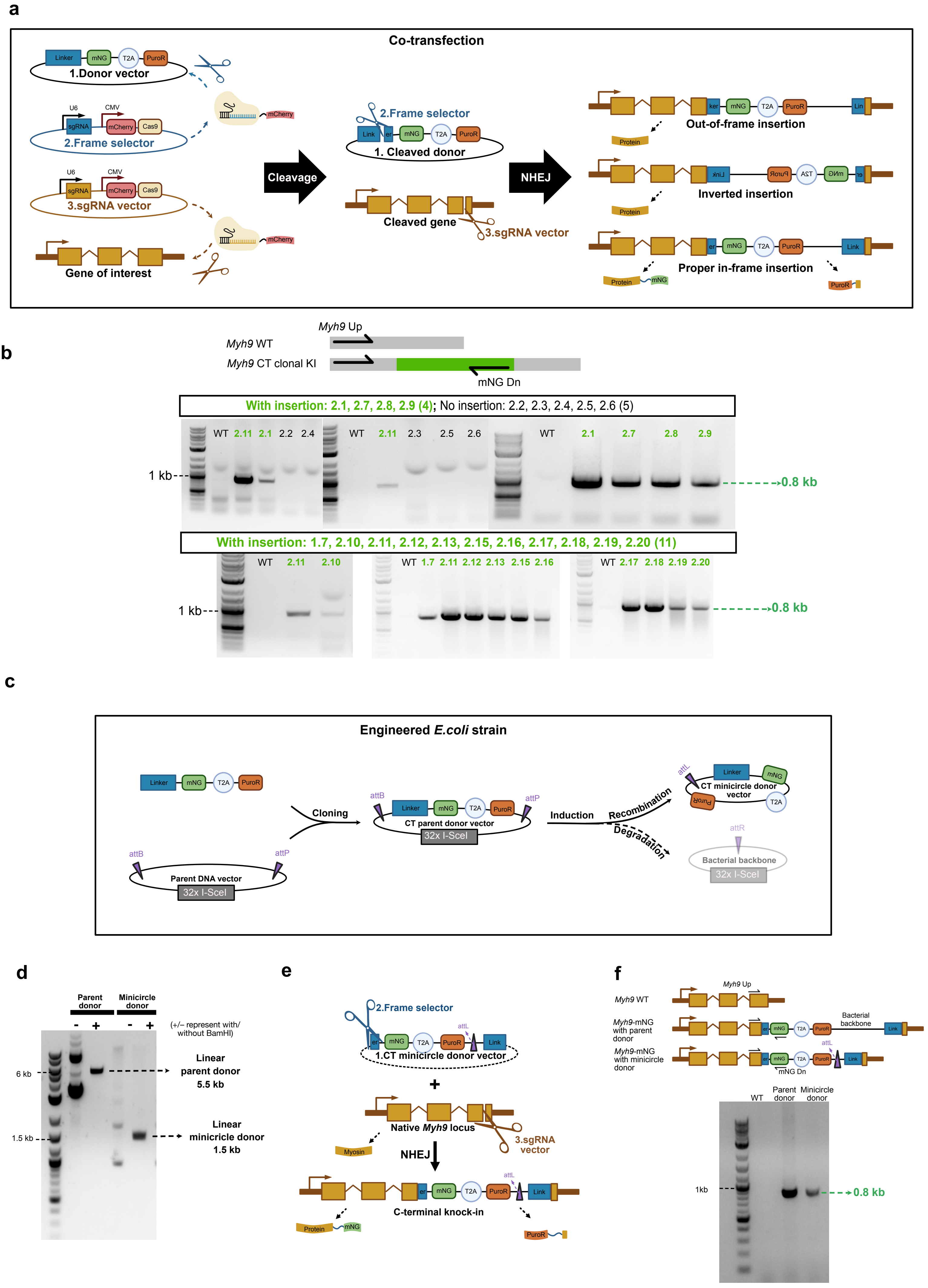

a
Co-transfection
Linker
mNG
T2A
PuroR
1.Donor vector
U6
CMV
sgRNA
mCherry
Cas9
2.Frame selector
3.sgRNA vector
Gene of interest
Cleavage
2.Frame selector
1. Cleaved donor
Cleaved gene
3.sgRNA vector
NHEJ
Out-of-frame insertion
Inverted insertion
Proper in-frame insertion
Protein
b
Myh9 Up
Myh9 WT
Myh9 CT clonal KI
mNG Dn
With insertion: 2.1, 2.7, 2.8, 2.9 (4); No insertion: 2.2, 2.3, 2.4, 2.5, 2.6 (5)
WT 2.11 2.1 2.2 2.4
WT 2.11 2.3 2.5 2.6
WT 2.1 2.7 2.8 2.9
1 kb
0.8 kb
With insertion: 1.7, 2.10, 2.11, 2.12, 2.13, 2.15, 2.16, 2.17, 2.18, 2.19, 2.20 (11)
WT 2.11 2.10
WT 1.7 2.11 2.12 2.13 2.15 2.16
WT 2.17 2.18 2.19 2.20
1 kb
0.8 kb
c
Engineered E.coli strain
Linker
mNG
T2A
PuroR
Cloning
attB
attP
Parent DNA vector
32x I-SceI
CT parent donor vector
32x I-SceI
Induction
Recombination
Degradation
attL
CT minicircle donor vector
attR
Bacterial backbone
32x I-SceI
d
Parent donor
Minicircle donor
- + - +
(+/– represent with/ without BamHI)
6 kb
1.5 kb
Linear parent donor 5.5 kb
Linear minicircle donor 1.5 kb
e
2.Frame selector
attL
1.CT minicircle donor vector
+
Native Myh9 locus
3.sgRNA vector
Myosin
NHEJ
C-terminal knock-in
Protein
mNG
PuroR
f
Myh9 Up
Myh9 WT
Myh9-mNG with parent donor
Bacterial backbone
Myh9-mNG with minicircle donor
attL
mNG Dn
WT
Parent donor
Minicircle donor
1kb
0.8 kb

**Figure S1: Adapting CRISPaint for C-terminal knock-ins in mESCs**
**S1a.** Illustration of the CRISPR tagging process. After co-transfection of three plasmids in cells, the target sgRNA vector cuts the desired gene and the frame selector cuts the linker region in the donor vector. The cleaved gene directly ligates with the linearized donor via non-homologous end joining (NHEJ). Only if the ligation is in-frame and in proper orientation between cleaved gene and linearized donor will the mNG-fused protein of interest and drug-resistance protein be produced. **S1b.** PCR genotyping of *Myh9*-mNG clonal cells. *Myh9* Up and mNG Dn primers amplified a specific band in *Myh9*-mNG cells compared with WT cells indicating a successful insertion. **S1c.** Illustration of C-terminal minicircle donor production from the C-terminal parent donor vector. **S1d.** DNA gel showing C-terminal minicircle donor with a correct size and a good purity. **S1e.** Illustration of C-terminal insertion by minicircle donor. The target sgRNA vector cuts the last exon of *Myh9* and the frame selector cuts the linker region in the minicircle donor. The cleaved *Myh9* directly ligates with the linearized donor via NHEJ. T2A separates the knock-in gene into two protein products. **S1f.** PCR genotyping of *Myh9*-mNG made using either parent or minicircle donor vector. In both conditions, *Myh9* Up and mNG Dn primers amplified a specific band compared to WT cells indicating a successful insertion.
Abbreviations: C-terminal (CT), mNeonGreen (mNG), Puromycin resistance gene (PuroR), the T2A peptide (T2A), Wildtype (WT), Knock-in (KI), attB and attP are recombination sites, attL and attR are byproducts of site-specific recombination.

### Figure S2

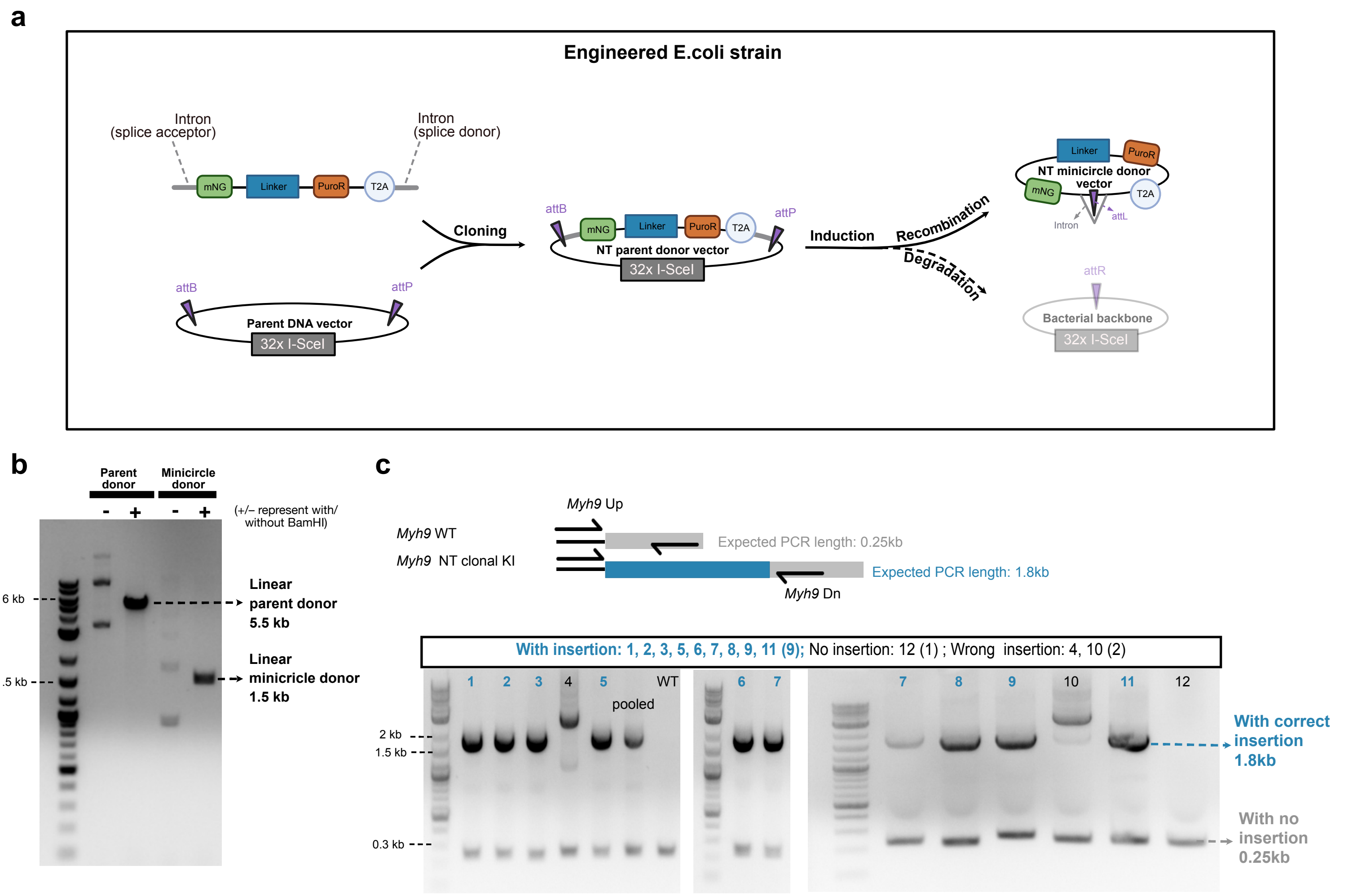


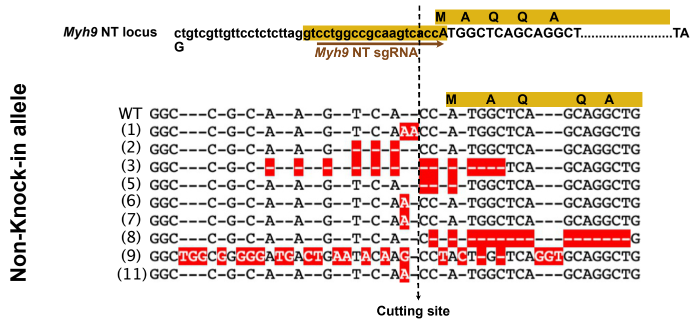


| | Knock-in allele at 5' junction | Knock-in allele at 3' junction | Non knock-in allele |
|---|---|---|---|
| WT | none | none | correct |
| 1 | -11nt | -24nt | +2nt |
| 2 | -5nt | -6nt | -3nt |
| 3 | 2nt mutation | -6nt | -13nt (missing start codon ) |
| 5 | -5nt | -6nt | -3nt |
| 6 | Correct | Correct | +1nt |
| 7 | -4nt | -3nt | +1nt |
| 8 | Correct | -36nt | -16nt (missing start codon ) |
| 9 | 5nt mutation | correct | +23nt (missing start codon ) |
| 11 | -4nt | -9nt | +1nt |

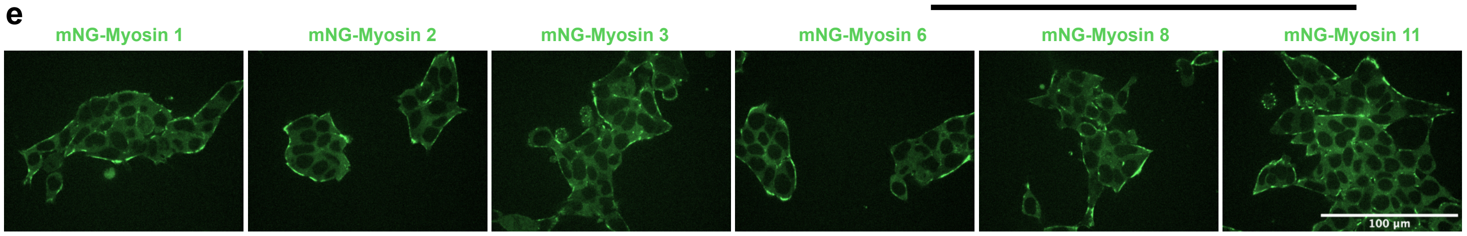


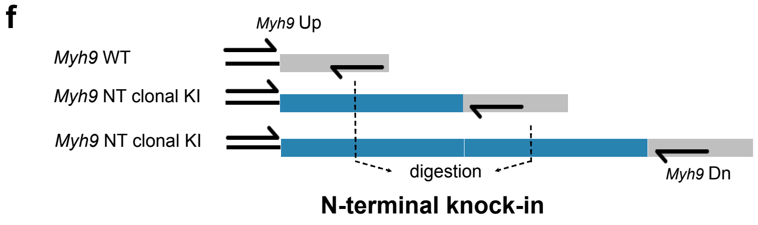


| | Allele1: Expected PCR length | Allele1: Expected digestion length | Allele2: Expected PCR length | Allele2: Expected digestion length |
|---|---|---|---|---|
| WT | 0.25 kb | 0.25 kb | 0.25 kb | 0.25 kb |
| One donor | 1.8 kb | 1 kb+0.8 kb | 0.25 kb | 0.25 kb |
| Two donor | 3.3 kb | 1k b+0.8 kb+1.5 kb | 0.25 kb | 0.25 kb |

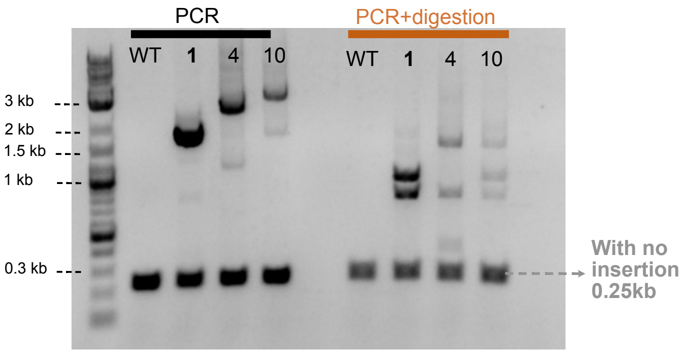

**Figure S2: Efficient knock-in at N-terminus of myosin**

**S2a.** Illustration of N-terminal minicircle donor production from N-terminal parent donor vector. **S2b.** DNA gel showing N-terminal minicircle donor with a correct size and a good purity. **S2c.** PCR genotyping of mNG-*Myh9* clonal cells. *Myh9* Up and Dn primers amplified a specific long band in mNG-*Myh9* cells compared with WT cells indicating a successful insertion. **S2d.** (Left) Alignment of Sanger sequencing results showing indels in the non-tagged alleles of clonal knock-in cell lines, indicating efficient cutting by the target sgRNA vector but less efficient tag insertion by NHEJ. (Right) Table summarizing sequencing results in picked clonal cells. **S2e.** Imaging of different clonal cells showing consistent expression and distribution of mNG-*Myh9.* Scale bar represents 100 μm. **S2f.** Digestion results showing the possibility of head-to-tail insertion with two minicircle donors into the same target site. Top left is a schematic graph for digestion in different conditions. Bottom left is a table with estimation of band size after digestion in different conditions. Right is the DNA gel of digestion result.

### Figure S3

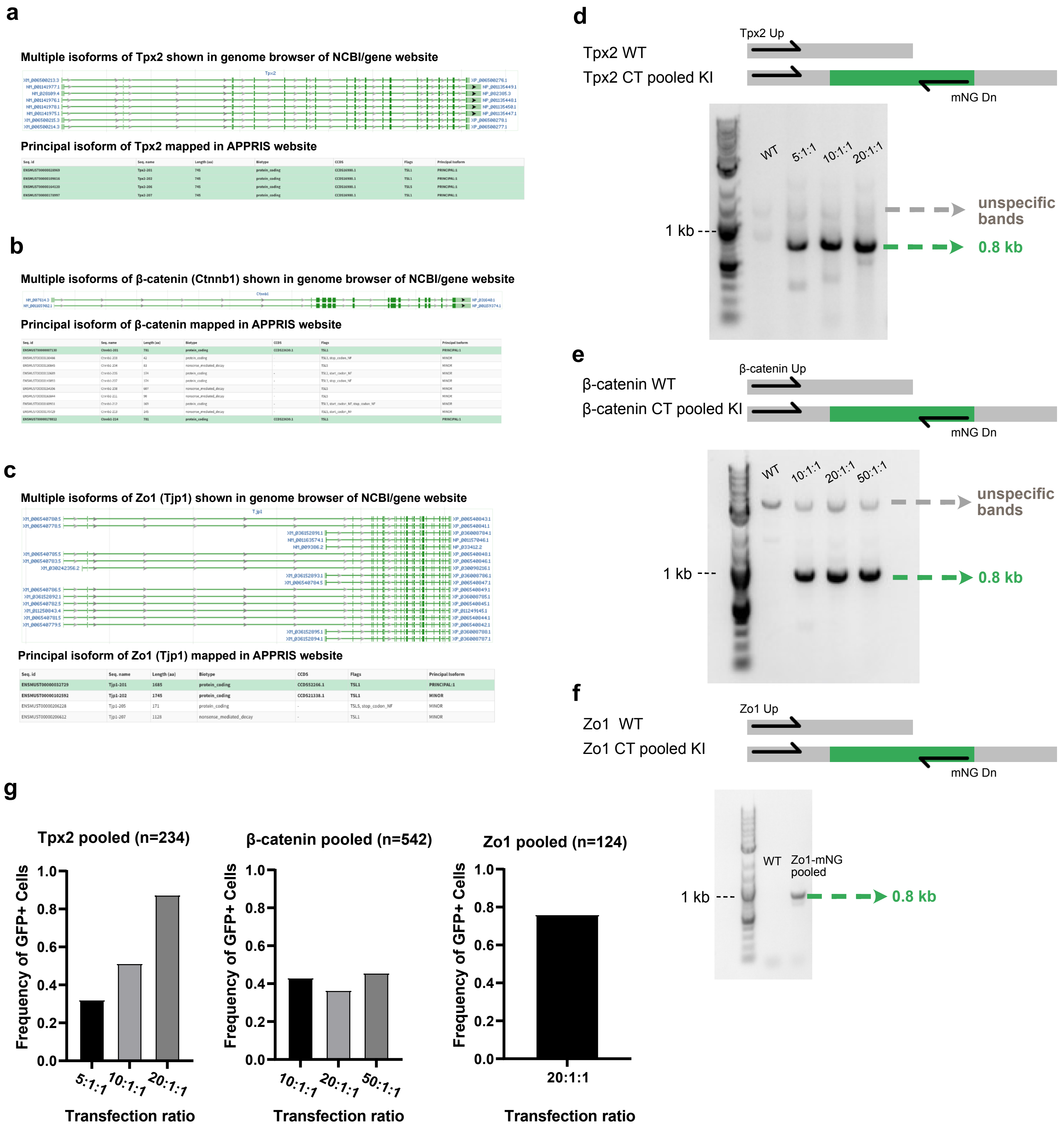


**Figure S3: Rapid transfection to imaging of pooled edited cells**
**S3a-b.** NCBI gene website was used to check gene structure and gene isoforms from the genome browser. The APPRIS website was used to identify the principal isoform. **S3d-f.** PCR genotyping of Tpx2-mNG, β-catenin-mNG, and Zo1-mNG pooled cells. All experiments showed a specific band amplified by a forward primer upstream of cutting site and mNG Dn primer, indicating successful insertions. **S3g.** The frequency of GFP+ cells was measured by microscopy in Tpx2-mNG, β-catenin-mNG, and Zo1-mNG pooled cells after transfection using different molar ratios of donor:sgRNA.

### Figure S4

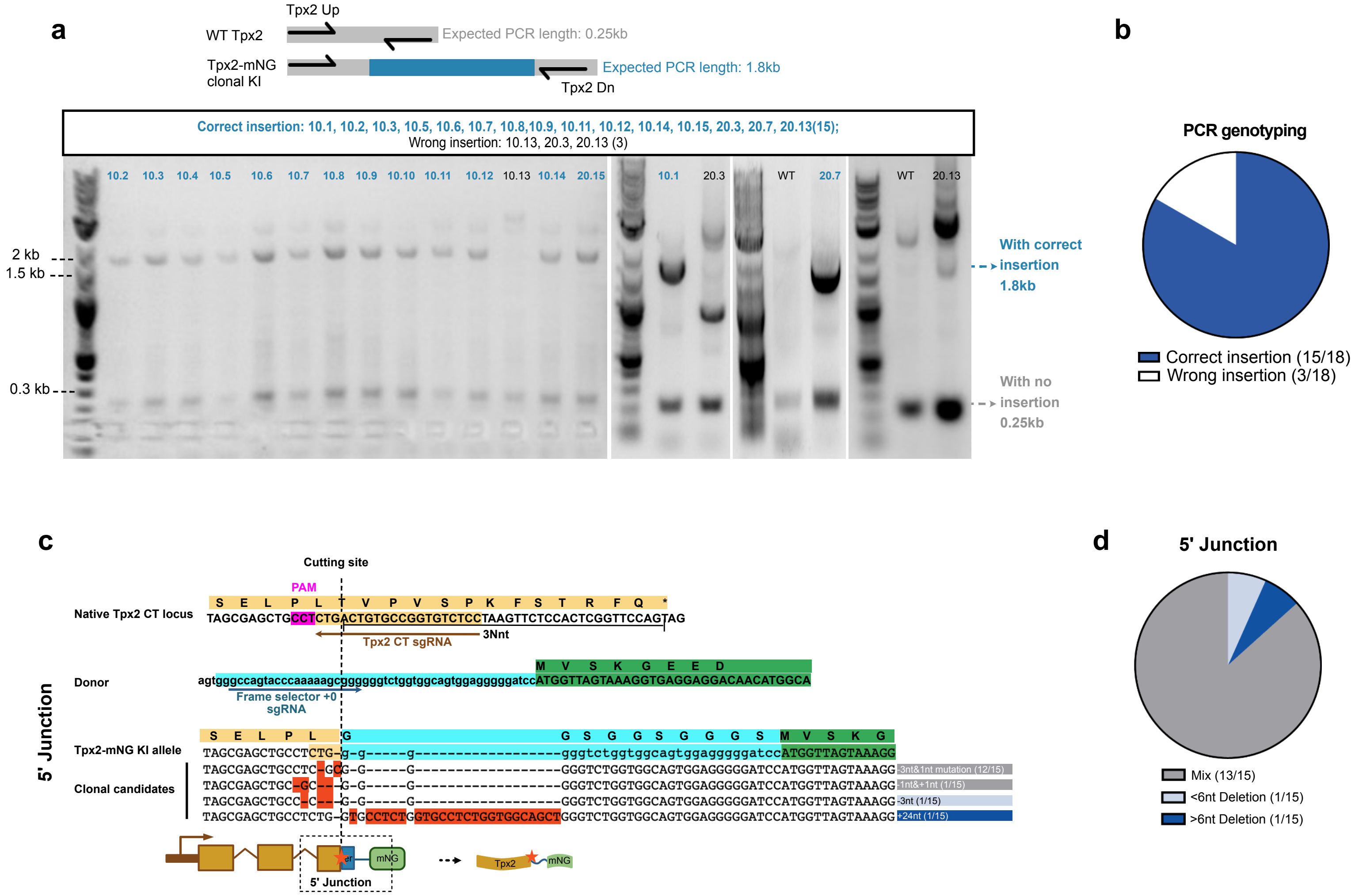


**Figure S4: Further analysis of Tpx2-mNG clonal cells**
**S4a.** PCR genotyping of Tpx2-mNG clonal cells. Tpx2 Up and Dn primers amplified a specific long band in Tpx2-mNG cells compared with WT cells indicating a successful insertion. **S4b.** Pie chart showing insertion efficiency in picked clonal cells. **S4c.** Sequence alignments showing the cleavage position of the target sgRNA in the target gene (top); the cleavage position of frame selector +0 in the donor vector (middle); and Sanger sequencing results in picked clonal cells (bottom). “-” in clonal sequences represents deletions. “-” in the reference sequence represents insertions. Highlighted nucleotides in clonal sequences indicate mutations. The red star in the cartoon indicates the locations of indels or mutations. **S4d.** Pie chart summarizing insertion results in picked clonal cells. Mix means a mixture of indels or/and mutation.

Abbreviations: C-terminal (CT), mNeonGreen (mNG), Puromycin resistance gene (PuroR), the T2A peptide (T2A), Wildtype (WT), Knock-in (KI), nucleotides (nt), 3Nnt where N represents the codon number.

### Figure S5

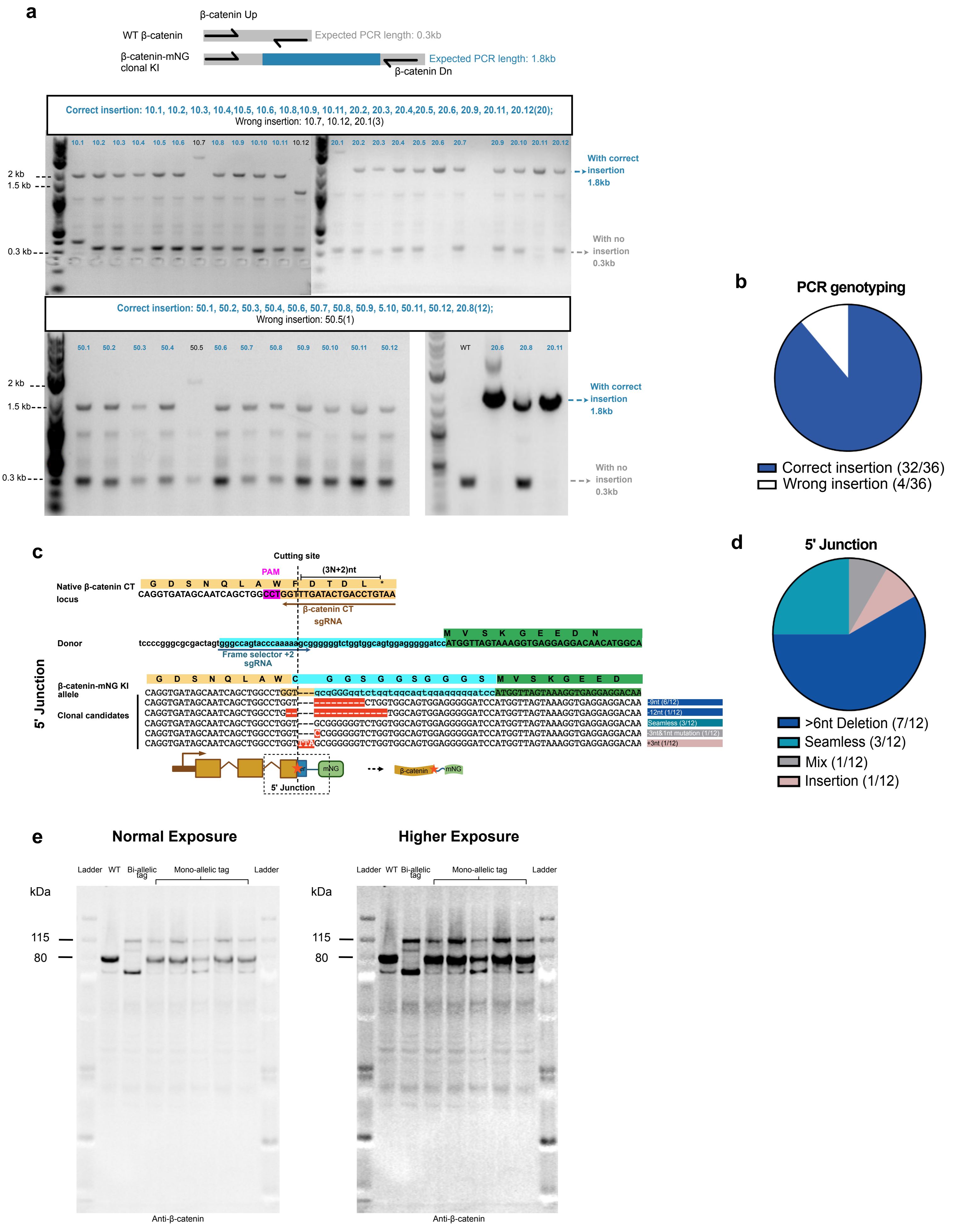
a
β-catenin Up
WT β-catenin
Expected PCR length: 0.3kb
β-catenin-mNG clonal KI
Expected PCR length: 1.8kb
β-catenin Dn
Correct insertion: 10.1, 10.2, 10.3, 10.4,10.5, 10.6, 10.8,10.9, 10.11, 20.2, 20.3, 20.4,20.5, 20.6, 20.9, 20.11, 20.12(20);
Wrong insertion: 10.7, 10.12, 20.1(3)
2 kb
1.5 kb
0.3 kb
With correct insertion 1.8kb
With no insertion 0.3kb
Correct insertion: 50.1, 50.2, 50.3, 50.4, 50.6, 50.7, 50.8, 50.9, 5.10, 50.11, 50.12, 20.8(12);
Wrong insertion: 50.5(1)
WT
b
PCR genotyping
Correct insertion (32/36)
Wrong insertion (4/36)
c
Cutting site
PAM
(3N+2)nt
Native β-catenin CT locus
CAGGTGATAGCAATCAGCTGGCCTGGTTTGATACTGACCTGTAA
β-catenin CT sgRNA
Donor
Frame selector +2 sgRNA
β-catenin-mNG KI allele
Clonal candidates
5' Junction
d
5' Junction
>6nt Deletion (7/12)
Seamless (3/12)
Mix (1/12)
Insertion (1/12)
e
Normal Exposure
Higher Exposure
Ladder
WT
Bi-allelic tag
Mono-allelic tag
kDa
115
80
Anti-β-catenin

**Figure S5: Further analysis of β-catenin-mNG clonal cells**

**S5a.** PCR genotyping of β-catenin-mNG clonal cells. β-catenin Up and Dn primers amplified a specific long band in β-catenin-mNG cells compared with WT cells indicating a successful insertion. **S5b.** Pie chart showing insertion efficiency in picked clonal cells. **S5c.** Sequence alignments showing the cleavage position of the target sgRNA in the target gene (top); the cleavage position of frame selector +2 in the donor vector (middle); and Sanger sequencing results in picked clonal cells (bottom). "-" in clonal sequences represents deletions. "-" in the reference sequence represents insertions. Highlighted nucleotides in clonal sequences indicate mutations. The red star in the cartoon indicates the locations of indels or mutations. **S5d.** Pie chart summarizing insertion results in picked clonal cells. Mix means a mixture of indels or/and mutation. S5e. Western blotting β-catenin-mNG clonal cells with mono-allelic or bi-allelic tag insertions. Left is Normal exposure and right is higher exposure.

Abbreviations: C-terminal (CT), mNeonGreen (mNG), Puromycin resistance gene (PuroR), the T2A peptide (T2A), Wildtype (WT), Knock-in (KI), nucleotides (nt), 3N+2nt where N represents the codon number.
