## Supplemental Protocol for "Efficient and rapid fluorescent protein knock-in with universal donors in mammalian stem cells"

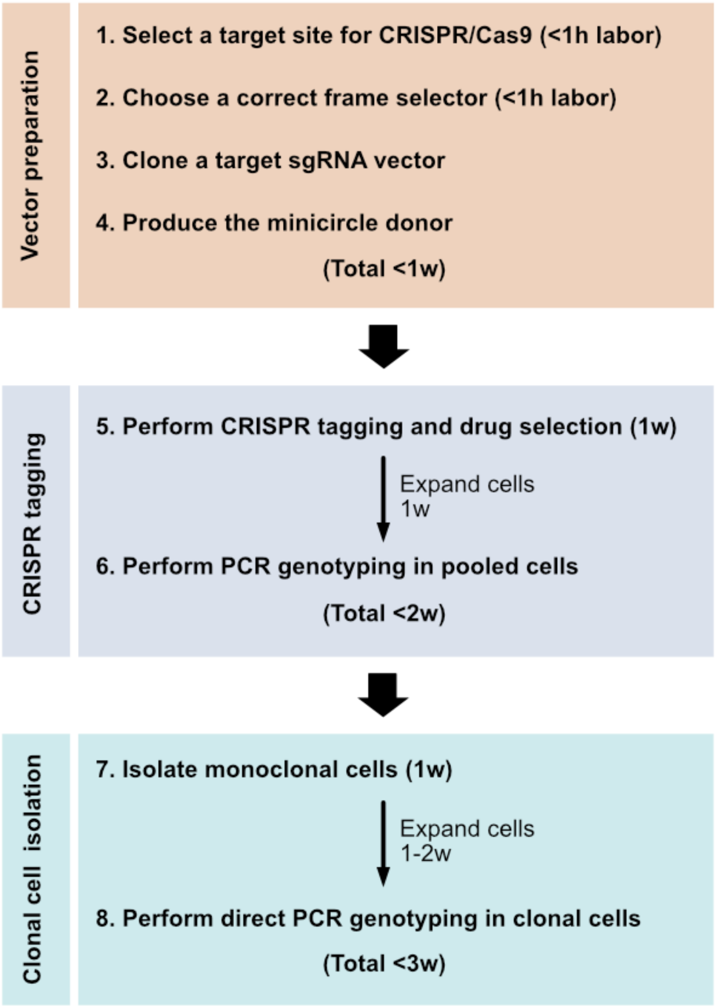

This protocol requires three plasmids: a fluorescent protein (FP) donor vector, a frame selector sgRNA vector and a target sgRNA vector. The target sgRNA vector is the only plasmid that needs to be cloned for the specific gene of interest; the other plasmids are premade and can be used ubiquitously.

### 1. Select a target site for CRISPR/Cas9 (<1h labor)

**Aim:** CRISPR tagging can utilize a single guide RNA (sgRNA) to target the gene of interest. To design a specific sgRNA, there are several considerations. First, mammalian cells express a wide range of alternatively spliced transcripts in a gene. Each transcript, or version of the gene, may require a different target site for CRISPR/Cas9. Therefore, the presence of multiple isoforms increases the complexity of designing sgRNA. Fortunately, it's been reported that most highly expressed protein-coding genes have a single dominant isoform[1]. Below is an example of Zo1 (Figure 1).

**a. Multiple isoforms of Zo1 (Tjp1) shown in genome browser of NCBI/gene website**

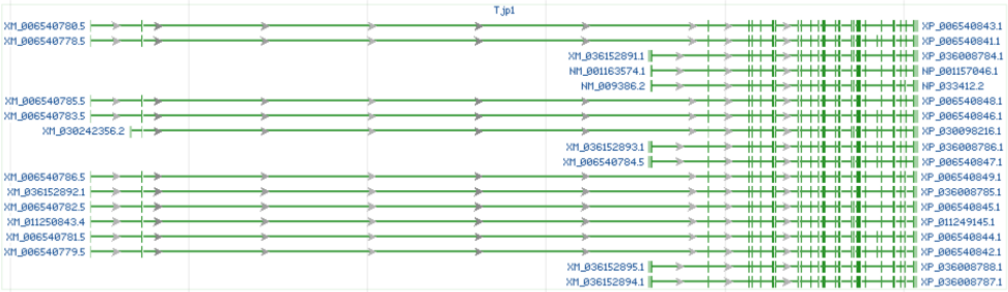

**b. Principal isoform of Zo1 (Tjp1) mapped in APPRIS website**

| Seq. id | Seq. name | Length (aa) | Biotype | CCDS | Flags | Principal Isoform |
|---|---|---|---|---|---|---|
| **ENSMUST00000032729** | **Tjp1-201** | **1685** | **protein_coding** | **CCDS52266.1** | **TSL1** | **PRINCIPAL:1** |
| **ENSMUST00000102592** | **Tjp1-202** | **1745** | **protein_coding** | **CCDS21338.1** | **TSL1** | **MINOR** |
| ENSMUST00000206228 | Tjp1-205 | 171 | protein_coding | - | TSL5, stop_codon_NF | MINOR |
| ENSMUST00000206612 | Tjp1-207 | 1128 | nonsense_mediated_decay | - | TSL1 | MINOR |

**Figure 1 Genome browser and principal isoform of Zo1**
**1a.** NCBI/gene website was used to check gene structure and gene isoforms from the genome browser. **2b.** APPRIS website was used to identify the principal isoform.

Determining the dominant isoform enables us to choose a Cas9 target site that is a reasonable choice for a first attempt at tagging a gene. It is always possible to choose other sites that target other isoforms later, if needed. To find the dominant isoform, we will use APPRIS[2], a web tool that flags the principal isoform based on protein structure, expression, and conserved function. Second, we will choose which end of the gene to tag. To avoid the perturbation of gene function by tagging, we use available information about the gene, from NCBI or other sources, to avoid functional motifs at the N- or C-terminus, where known. Third, the CHOPCHOP database can be used for selecting target site for Cas9. We aim to pick a target site that is close to the region of interest with high specificity and reasonable efficiency.

**Useful websites:**

1. APPRIS: https://appris.bioinfo.cnio.es/#/seeker
2. NCBI/gene: https://www.ncbi.nlm.nih.gov/gene
3. CHOPCHOP: https://chopchop.cbu.uib.no/

**Protocol:**

1. Identify the dominant isoform for the gene of interest. Use the APPRIS (see the link above) by inputting the gene name and species.
2. Decide which terminus of the protein of interest you want to tag. Based on NCBI or other useful databases, verify the structure or sequence conservation of the gene. Choose the end that is less likely to perturb protein function by avoiding known, conserved functional sequences.
3. Select a target site to cleave the near the end of the gene using CHOPCHOP. There are several considerations that dictate the choice of target sites. First, aim for a cut site that is close to the start codon or right before the stop codon. When the cut site is close to the N- or C-terminus, it's less likely to disrupt protein expression and function. Second, determine target sites with the fewest possible off-targets. Third, select a target site with a higher predicted cutting efficiency. Of note, specificity of a target site is more important than efficiency to obtain minimal off-targets because editing via the NHEJ pathway is efficient.
4. Order 20nt target oligo and clone the target oligo into our modified target sgRNA vector (Cloning details are in section 3).

#### 2. Choose a correct frame selector (<1h labor)

**Aim:** For CRISPR tagging at either C- or N-terminal (CT or NT) site, we provide three premade frame selectors (CT: +0, +1, +2; NT: -0, -1, -2). Among them, frame selector -2 is a novel design to facilitate N-terminal insertion. The CMV promoter is inactive in mouse embryonic stem cells (mESCs)[3], so we have adapted frame selectors for mESCs by replacing the CMV promoter with the EF1α promoter to express Cas9. To favor in-frame integration of the cleaved genome and donor sequence, it's important to select a correct frame selector based on where the chosen sgRNA cuts the gene of interest (Table 1). Because the target sgRNA vector may cleave the genome at one of three possible positions related to the reading frame, the frame selector can also cleave donor vectors at one of three adjacent nucleotide positions from the cutting site. Thus, using the correct frame selector favors in-frame insertion.

| Insertion terminus | Frame selectors | Cas9 target within donor vectors | Cleavage to stop codon |
|---|---|---|---|
| CT | pDD425 Frame +0 | Gccagtacccaaaaagcggg | 3N |
| | pDD426 Frame +1/–0 | Ggccagtacccaaaaagcgg | 3N+1 |
| | pDD427 Frame +2/–1 | Gggccagtacccaaaaagcg | 3N+2 |
| NT | pDD426 Frame +1/–0 | Ggccagtacccaaaaagcgg | 3N |
| | pDD427 Frame +2/–1 | Gggccagtacccaaaaagcg | 3N+1 |
| | pYS19 Frame –2 | Cgggccagtacccaaaaagc | 3N+2 |

**Table 1: Frame selector information**

For either C- or N- terminal insertion, there are three frame selectors. Different frame selectors use different Cas9 targets to cut one of three adjacent positions on linker region of donor vectors, which helps to realign the reading frame. The choice of a correct frame selector is based on the distance from the cutting site to stop codon on genome. There are three possible distances: 3N, 3N+1, 3N+2 nucleotides, where N represents the number of codons. The distance can be estimated by target sgRNA vector which cuts the gene of interest. Of note, N terminal insertion can also use stop codon as reference regardless of whether the target site cuts before or after start codon of gene (Make sure not to count introns any codon number).
Abbreviations: C-terminal (CT), N-terminal (NT).

Taking the C-terminal insertion of *Myh9* as an example, mNG will be fused with the 3' end of myosin, so in-frame insertion at junction upstream of the cut site is important. The chosen sgRNA cuts 26 nucleotides (3N+2, where N is the number of codons – 8 in this case) away from the stop codon. 2 nucleotides will be lost at the junction upstream of the cut site relative to the reading frame. Frame selector +2 will favor in-frame insertions by providing 2 extra nucleotides in the linker region before the coding sequence of mNG. Thus, Frame selector +2 is recommended. (Figure 2)

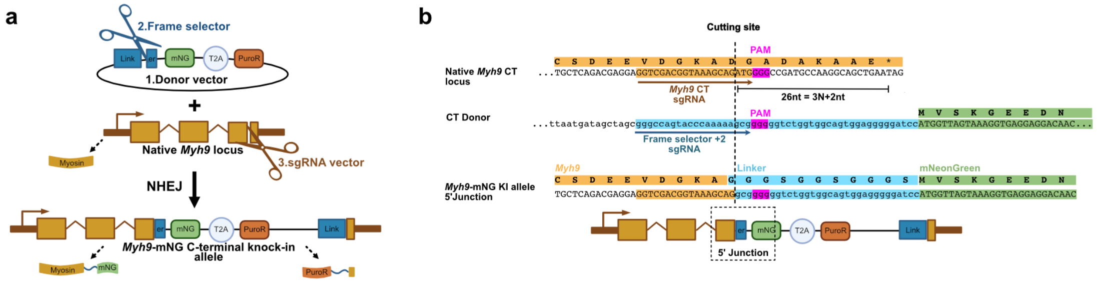

**Figure 2 The rule of selecting a correct frame selector for C-terminal insertion of *Myh9***
**2a**. Illustration of C-terminal insertion via non-homologous end joining (NHEJ). The target sgRNA vector cuts the last exon of *Myh9* and the frame selector cuts the linker region in the donor vector. The cleaved *Myh9* directly ligates with the linearized donor via NHEJ. T2A separates the knock-in gene into two protein products. **2b.** Sequence alignments showing the cleavage position of the target sgRNA in the target gene (top); the cleavage position of frame selector +2 in the donor vector (middle); and the expected product (bottom). *Myh9* CT sgRNA cuts 26 nucleotides away from the stop codon (3N+2 nt, where N represents the codon number). Frame selector +2 realigns cleaved *Myh9* by offering 2 extra nucleotides to favor the in-frame insertion. Abbreviations: C-terminal (CT), mNeonGreen (mNG), Puromycin resistance (PuroR), theT2A peptide (T2A), Knock-in (KI).

Another example is N-terminal tagging of *Myh9*. To ensure gene expression of N-terminal tags, the insertion must be in-frame at both junctions of the cleavage site. The chosen sgRNA gives rise to 2nt from the cutting site to the stop codon (3N+2, where N is the number of codons), resulting in a loss of 2 nucleotides at 5' junction and a gain of 2 nucleotides at 3' junction relative to the reading frame. Frame selector -2 realigns the reading frame by providing 2 extra nucleotides for the 5' junction and 1 extra nucleotide for the 3' junction in linker region of the N-terminal donor. Thus, Frame selector -2 is recommended. (Figure 3)

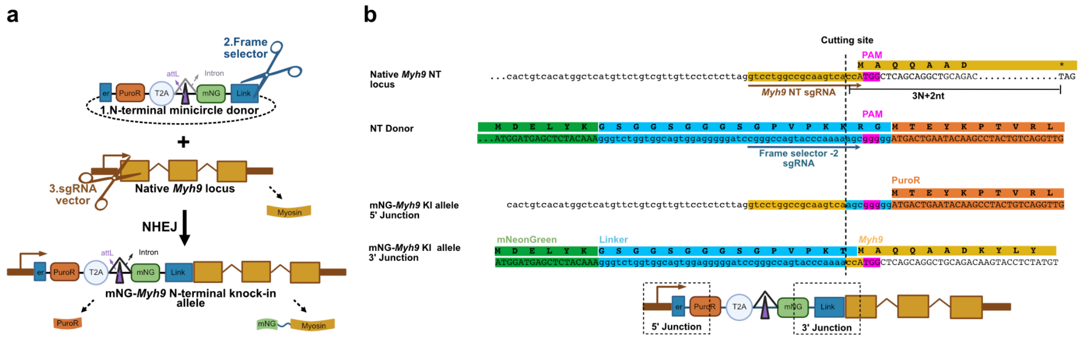

**Figure 3 The rule of selecting a correct frame selector for N-terminal insertion of *Myh9***
**3a**. Illustration of N-terminal insertion via NHEJ. The target sgRNA vector cuts the 5' UTR of *Myh9* and the frame selector cuts the linker region in the N-terminal minicircle donor. The cleaved *Myh9* directly ligates with the linearized donor via NHEJ. T2A separates the knock-in gene into two protein products. **3b.** Sequence alignments showing the cleavage position of the target

sgRNA in the target gene (top); the cleavage position of frame selector -2 in the donor vector (middle); and the expected product at both junctions (bottom). *Myh9* NT sgRNA cuts 3N+2nt away from the stop codon (where N represents the codon number). Frame selector -2 compensates cleaved *Myh9* with extra nucleotides to favor the in-frame insertion.

#### 3. Clone a target sgRNA vector (<1w)

**Aim**: To cleave the gene of interest, the chosen target sequence is inserted into the target Cas9/sgRNA vector (Figure 4).

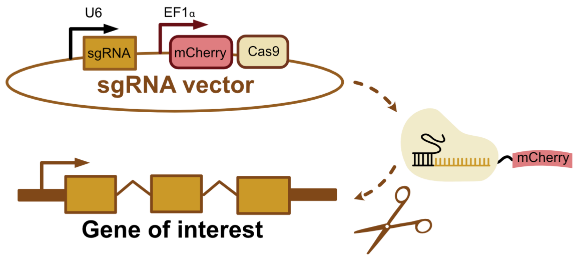

**Figure 4 The structure and function of the target sgRNA vector.**
The target sgRNA vector contains U6 promoter to express an sgRNA (target and Scaffold RNA), and EF1α promoter to express mCherry fused Cas9. After expression, Cas9 is guided by the sgRNA to execute cleavage on the gene of interest.

To clone the target sgRNA vector, pDD428 vector can be used as a template. The sgRNA consists of two parts: the target site (protospacer), a 20-nucleotide sequence complementary to the gene of interest, and a scaffold RNA, which serves as a binding scaffold for the Cas9. The idea of cloning a new sgRNA vector is to replace the original *Myh9* target site in pDD428 with a new target site. Of note, U6 requires a G residue as the first base of the sgRNA sequence to initiate transcription. We appended an extra G residue to the 5' end of the target site since extensions of the target site beyond 20 bp do not affect cleavage activity[4]. Therefore, a user does not need to consider the first nucleotide when selecting a target site.

There are two alternative cloning strategies. One is Gibson assembly, the other is KLD cloning. The following diagrams summarize the two cloning protocols (Figure 5). The advantage of KLD cloning is to save labor. However, PCR amplification of pDD428 vector (over 10 kb) can be difficult, which makes KLD cloning not as efficient as Gibson assembly. Thus, we prefer Gibson assembly over KLD cloning in most cases.

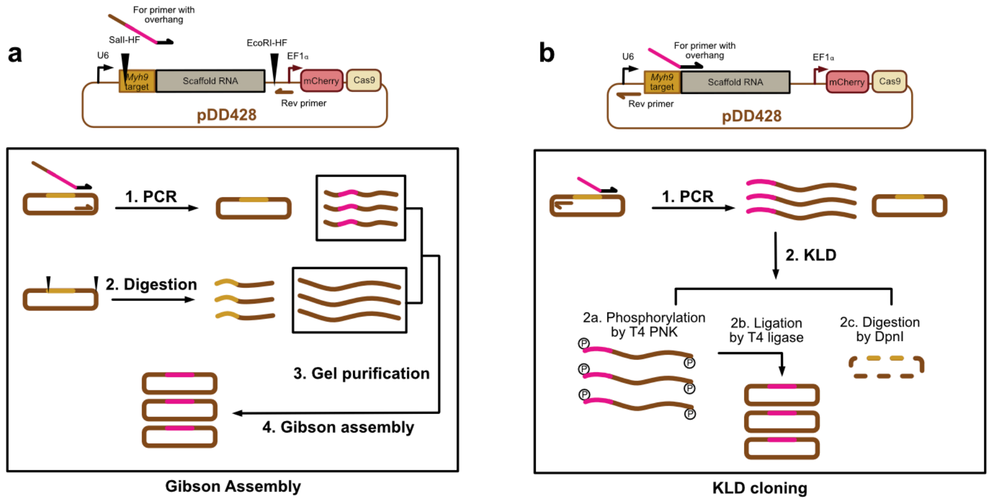

**Figure 5 Illustration of target Cas9/sgRNA vector cloning strategies**
pDD428 is the template vector containing a U6 promoter to express *Myh9* sgRNA (*Myh9* target and Scaffold RNA), and an EF1α promoter to express mCherry-Cas9. The idea of cloning is to replace a new target (magenta) with an *Myh9* target (yellow) in pDD428. **5a**. Gibson assembly requires a PCR and a digestion reaction. PCR needs a forward primer with overhang (20nt new target plus 20nt Gibson overlap) and a universal reverse primer at the EF1α promoter. Then Gibson assembly can be performed by using purified PCR insert and the parent vector, which is prepared by digesting with SalI-HF and EcoRI-HF. **5b.** KLD cloning requires a PCR via a forward primer with overhang (20nt new target) and a universal reverse primer at U6 promoter. The KLD reaction contains kinase, ligase and DpnI enzymes which self-ligate the PCR product to produce a new vector.

#### Materials:

1. Q5 High-Fidelity DNA Polymerase (NEB M0491)
2. Digestion enzymes: SalI-HF (NEB R3138S), EcoRI-HF (NEB R3101S)
3. Zymoclean Gel DNA Recovery Kits (Zymo D4007/D4008) or equivalent.
4. NEBuilder HiFi DNA Assembly Master Mix (NEB E2621S)
5. T4 polynucleotide kinase (NEB M0201S)
6. T4 DNA ligase (NEB M0202S)
7. T4 DNA ligase buffer 10x (NEB B0202S)
8. DpnI (NEB R0176S)
9. Homemade KLD 1x buffer: 1 µL of T4 DNA ligase buffer 10x (with ATP inside already) + 9 µL of Nuclease-free water
10. Homemade KLD enzyme mix: 16.7 µL PNK + 1 µL T4 ligase + 8.3 µL DpnI
11. DH10B competent *E. coli* (NEB C3019HVIAL)
12. NEB 10-beta/Stable Outgrowth Medium (NEB B9035SVIAL)
13. LB Agar plate with 100 µg/mL Ampicillin.

#### Gibson assembly protocol:

1. Using 5 ng of pDD428 as template, perform Q5 PCR to amplify the sgRNA region pDD428 and replace the original *Myh9* target site. The Forward primer including the new target site is 5'-gtggaaaggacgaaacaccGNNNNNNNNNNNNNNNNNNNNGTTTTAGAGCTAGAAATAGC-3', and the Reverse primer is 5'-GCGATGATAACGCGTATATC-3', which is universal for cloning different sgRNA vectors.
2. Digest 1 µg of pDD428 with SalI-HF and EcoRI-HF.

3. Run a gel with the PCR and digestion products. The expected length of PCR is 0.16 kb, the expected length of digested vector is 10.5 kb.
4. Purify 0.16 kb PCR insert and 10.5 kb digested vector by gel extraction.
5. To perform Gibson assembly, take 1 µL of purified PCR insert + 1 µL of purified digestion vector + 8 µL of Nuclease-free water + 10 µL of NEBuilder HiFi DNA Assembly Master Mix, incubate at 50ºC for 1 h.
6. Take 2-5 µL of Gibson assembly product and transform into DH10B competent *E. coli*.
7. Pick at least 6 candidates for screening. Grown overnight cultures and purify plasmid DNA.
8. Use a forward primer of U6 promoter 5'-GACTATCATATGCTTACCGT-3' to screen correct sgRNA cloning by Sanger sequencing.

**KLD cloning protocol:**

1. Using 5 ng of pDD428 as template, perform Q5 PCR to amplify pDD428 and replace the original *Myh9* target site. The Forward primer including the new target site is 5'-NNNNNNNNNNNNNNNNNNNNGTTTTAGAGCTAGAAATAGCAAGT-3', the Reverse primer is 5'-Cggtgtttcgtcctttc-3', which is universal for cloning different sgRNA vectors, the expected length of PCR is 10.6 kb.
2. To perform KLD reaction, mix 1 µL of unpurified PCR product + 8.5 µL of KLD buffer + 0.5 µL of KLD enzyme mix, incubate 10 min at room temperature, then 20 min at 37°C.
3. Take 2-5 µL of KLD product and transform into DH10B competent *E. coli*.
4. Pick at least 6 candidate clones for screening. Grow overnight cultures and purify plasmid DNA.
5. Use a forward primer of U6 promoter 5'-GACTATCATATGCTTACCGT-3' to screen correct sgRNA cloning by Sanger sequencing.

#### 4. Produce the minicircle donor (<1w)

**Aim:** We generated parent donor vectors for either C- or N-terminal insertion. These donors can be made into "minicircle donors" that lack bacterial sequences from the plasmid backbone; in this way, only the desired sequences (the fluorescent protein and drug selection marker) are inserted into the gene of interest. For N-terminal tags, the use of a minicircle donor is required to avoid disrupting expression of the gene of interest. For C-terminal tags, a minicircle donor is optional but is recommended to avoid disrupting the gene's 3'UTR. We provide 4 premade parent donor vectors which were transformed in a special *E. coli*, called ZYCY10P3S2T *E. coli*[5]. The genome of the *E. coli* has been modified to facilitate minicircle donor production. To make minicircle donors, ZYCY10P3S2T *E. coli* carrying either C- or N- terminal parent donor needs to be grown until the proper OD value is achieved, then induced by induction buffer. (Figure 6)

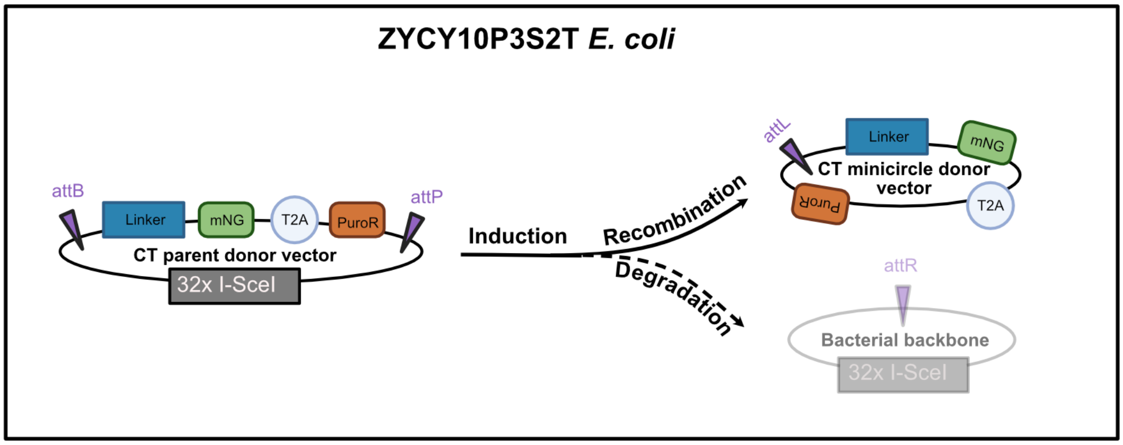

**Figure 6 Illustration of C-terminal minicircle donor production**
An engineered *E. coli* called ZYCY10P3S2T carries a C-terminal parent donor vector, in which attB-attP recombination sites flank the Linker–mNG–T2A–PuroR cassette. Through induction, attB/attP recombination circularizes into minicircle donor and the bacterial backbone is eliminated due to degradation mediated by 32 I-SceIs sites.

#### Materials:

1. LB Agar plate with 50 µg/mL Kanamycin
2. LB+Kan: LB containing 50 µg/mL Kanamycin.
3. TB+Kan: mix 47.6 g of Terrific Broth Powder (VWR J869-500G) and 4 mL of glycerol in 1 L of purified water, autoclave, and allow the solution to cool down before containing 50 µg/mL Kanamycin
4. 1N sodium hydroxide (NaOH): dissolve 40 g of NaOH in milli-Q water to make volume 1 L.
5. 20% L-arabinose: dissolve 2 g of L-Arabinose in milli-Q water to make volume 10 mL, sterilize it with a 0.22 µm filter. (Thermo Fisher AC104980250)
6. Induction buffer: mix 400 mL of fresh LB, 16 mL of 1N NaOH and 0.4 mL 20% L-arabinose.
7. QIAprep Spin Miniprep Kit (Qiagen 27104)
8. Vacufuge Concentrator (Eppendorf Mfr. No.022820109)
9. Whole plasmid sequencing: https://www.plasmidsaurus.com/

#### Protocol:

1. On d1, streak the parent donor from the glycerol stock on LB Agar plate with 50 µg/mL Kanamycin. Grow overnight at 37ºC.
2. On d2, inoculate four single colonies into four separate tubes containing 2 mL of LB+Kan. Incubate at 30°C shaking at 250 rpm for 4 h, test their OD values, choose one with a highest OD value, and store it at 4ºC overnight.
3. On d3, transfer 1 mL from the chosen liquid culture in a 500 mL flask with 50 mL of TB+Kan, incubate at 30°C shaking at 250 rpm for 4-6 h. Start to test OD value after 4h then test every half hour until the OD value is between 4 and 6. Add 50 mL of induction buffer to 50 mL of TB+Kan liquid culture whose OD has been checked, incubate the mixture at 30°C shaking at 250 rpm for 3h, then increase temperature to 37°C for 1h. (Of note, the ratio of flask size to culture volume at 5:1 (vol:vol) will ensure proper aeration for good bacterial growth.)

4. Collect and aliquot 100 mL induced E. coli into 3 mL of per tube.
5. To store the induced E. coli for future miniprep, spin down the aliquoted bacteria, remove supernatant and freeze in -20°C.
6. To extract minicircle donors, use the QIAprep Spin Miniprep Kit. Before miniprep, thaw one aliquot of induced E. coli at 4°C and resuspend the pellet with 500 μL of P1 buffer. Then lyse it with 500 μL of P2 buffer for 5 min and mix it with 700 μL of N3 buffer, centrifuge, and transfer supernatant to one QIAprep spin column. Wash the spin column by 500 μL of PB buffer and 750 μL of PE buffer. Finally, elute minicircle donors with 50 μL of Nuclease-free water.
7. Check the quality of the minicircle donor by linearizing the donor with BamHI and performing gel electrophoresis.
8. To verify minicircle donors, sequence whole plasmid by plasmidsaurus.
9. Concentrate minicircle donor using Vacufuge Concentrator.

**Note:**

1. To ensure yield and quality of minicircle donors, we recommend using 1 column to extract 3 mL of induced culture and using twice volume of P1, P2 and N3 buffer per column.
2. A 3 mL of induced *E. coli* can yield around 2.5 μg of minicircle donors (50 ng/μL in 50 μL), which can be used in around 6 co-transfections (364 ng minicircle donor will be used per co-transfection).
3. To obtain the optimum transfection efficiency, minicircle donors are concentrated to around 500 ng/μL.
4. Digestion (for example BamHI) and electrophoresis are recommended to verify if size is correct and if there is any contamination from bacterial genomic DNA or parent donor vectors. The left is a good example of minicircle donor extraction, the right is an example of minicircle donor with bacterial DNA contamination. (Figure 7)
5. Our parent donor vectors have been transformed in ZYCY10P3S2T *E. coli* which can be directly used for minicircle donor production. If there is a need of cloning another fluorescent gene, it requires a transformation in ZYCY10P3S2T *E. coli* before minicircle donor production.

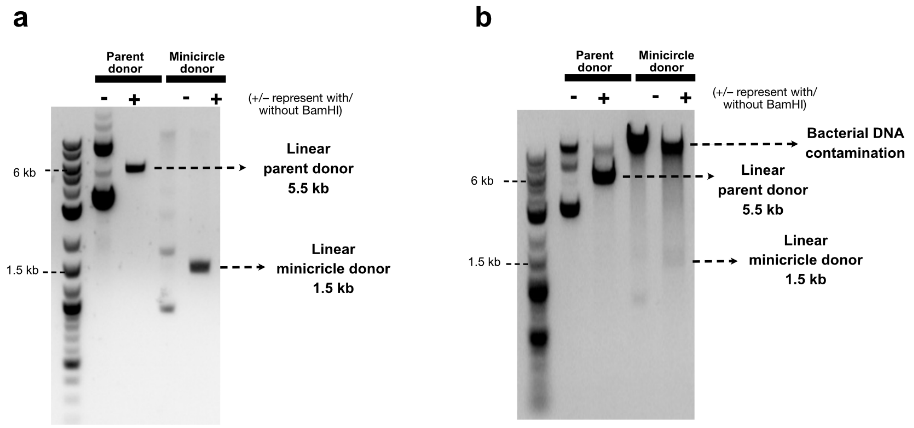

**Figure 7 Two examples of minicircle donor purification**
Parent donor is around 5.5 kb, minicircle donor is around 1.5 kb. BamHI is a restriction digestion enzyme that has one cutting site in either parent or minicircle donor. **7a**. A good example of minicircle donor extraction with dark and correct size of

minicircle donor as well as dim or no bacterial DNA. **7b.** A bad example of minicircle donor extraction with dark bacterial DNA but dim minicircle donor.

### 5. Perform CRISPR tagging and drug selection (1-2w)

**Aim:** To achieve fluorescent protein knock-in, we transfect three plasmids (an FP donor vector, a frame selector sgRNA vector and a target sgRNA vector) into mESCs using Lipofectamine 2000. The donor vector contains a FP gene fused with a flexible linker, a self-cleaving T2A peptide, and a drug resistance gene. A pre-made frame selector plasmid expresses Cas9 fused with mCherry and an sgRNA that cleaves the linker region of the donor vector at a specific position, so that it allows the linearized donor to compensate the reading frame of cleaved genome accordingly. A second Cas9/sgRNA vector cleaves the gene of interest. Of note, the donor vector does not contain its own promoter, so the FP and drug resistance sequences can only be expressed if the linearized donor is ligated to the cleaved gene in-frame and in the proper orientation.

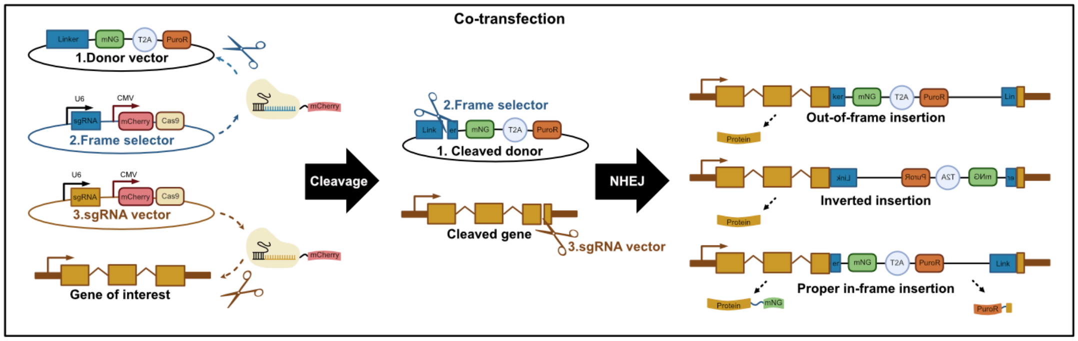

**Figure 8 Schematic process of CRISPR tagging via NHEJ**
After co-transfection of three plasmids in cells, target sgRNA vector cuts desired gene and Frame selector cuts linker region in donor vector. The cleaved gene directly ligates with linearized donor via NHEJ. Only if the ligation is in-frame and in proper orientation between cleaved gene and linearized donor, it will produce mNG fused protein of interest and drug-resistant protein.

We tested transfections with different [donor vector] : [frame selector] : [target sgRNA] plasmid molar ratios, ranging from 5:1:1 to 50:1:1. Although we didn't see a clear pattern that would indicate which ratio works best, they all can achieve successful knock-in. As a starting point, we recommend using 20:1:1.

**Materials:**

1. Gelatin-coated 12-well plate
2. mESCs growing in gelatin-coated 10cm plates (WT mESCs for mono-color tagging, tagged mESCs for dual-color tagging)
3. Accutase (Sigma SCR005)
4. Growth medium 2i/LIF: Commercial N2B27(ESGRO Complete Basal Medium, Sigma SF002-500) or Homemade N2B27[6] complemented with 3 μM of GSK3β inhibitor (Sigma SML1046), 1 μM of MEK inhibitor (Sigma PZ0162) and 100 U/mL of Leukemia Inhibitory Factor (Sigma ESG1106)
5. Wash buffer: 500 mL of DMEM/F12 (Sigma D6421) + 8 mL of bovine serum albumin fraction V (Thermo Fisher 15260037)
6. Trypan blue (Thermo Fisher 15250061)
7. Cell counter (Invitrogen AMQAF1000)

8. Plasmids: an FP donor vector, a frame selector sgRNA vector and a target sgRNA vector. To obtain optimum transfection and insertion efficiency, the starting concentration of each plasmid should be ≥500 ng/μL.
9. Lipofectamine 2000 (Thermo Fisher 11668027)
10. Opti-MEM I Reduced Serum Medium (Gibco 31985062)
11. Drug selection: 1-2 μg/mL Puromycin (Thermo Fisher A1113803) or 100-200 μg/mL Hygromycin (Thermo Fisher 10687010) in 2i/LIF

**Protocol:**

1. On d0, warm wash buffer, 2i/LIF and gelatin-coated 12-well plate.
2. Remove 2i/LIF from cells in gelatin-coated 10 cm plates, add 3.5 mL of Accutase and incubate for 5min at 37 °C, pipet up and down to dissociate mESCs into single cell suspension, and neutralize Accutase by adding 10.5 mL of wash buffer. Pellet cells for 5 min at 1000 rpm, remove supernatant, resuspend cells with 1 mL of 2i/LIF, and count cell number.
3. Dilute DNA with Opti-MEM Medium to a final volume of 50 μL. For a [donor vector] : [frame selector] : [target sgRNA] molar ratio of 20:1:1, use 364 ng minicircle donor,130 ng sgRNA vector, and 130 ng frame selector. Use 50 μL of Opti-MEM Medium with no DNA as a negative control.
4. Dilute 1.25 μL of Lipofectamine 2000 to a final volume of 50 μL.
5. Mix the 50 μL of DNA mixture or Opti-MEM negative control with the 50 μL of Lipofectamine 2000 mixture, and incubate at room temperature for 5 min. Then transfer the 100 μL of DNA/ Lipofectamine mixture or negative control mixture into two different wells of the gelatin-coated 12-well plate.
6. Add 5 X 10^5 single cell suspension per well based on the cell counting and add 2i/LIF to a final volume of 1 mL.
7. On d1, replace transfection mixture with 2-3 mL of fresh 2i/LIF and check mCherry-Cas9 expression using a fluorescence microscope.
8. On d2, perform drug selection (1-2 μg/mL Puromycin or 100-200 μg/mL Hygromycin in 2i/LIF depending on which donor vector has been used).
9. To remove floating dead cells, gently use wash buffer to wash cells for a couple of times but do not disturb adherent live cells. Replace with fresh drug-containing medium daily on d3-4.
10. On d5, replace drug with 2i/LIF to expand survival cells.
11. Change media every 2 days until cell number is enough for imaging, genomic DNA extraction, and subculture.

**Notes:**

1. Transfection of mESCs is reported to be more efficient by using single cell suspension and a [DNA]/[Lipofectamine 2000] ratio of 1: 2[7].
2. To achieve a [DNA] : [Lipofectamine 2000] of 1: 2, the total amount of DNA (364 ng minicircle donor +130 ng sgRNA vector +130 ng frame selector) is 0.624 μg, and the amount of Lipofectamine 2000 is 1.25 μL.
3. We typically perform transfections in triplicate, especially when targeting a new gene for the first time.
4. On d1, red fluorescent signal can be observed due to mCherry-Cas9 expression which indicates successful transfection of the frame selector and target sgRNA plasmids.
5. The goal of drug treatment is to enrich putatively edited cells. The length of drug treatment varies depending on the target gene, because the expression of the drug resistance marker relies on the gene's native promoter which can

be expressed at varying levels. The length of drug treatment is usually 2-3 days, but the length can be shortened when there are very few colonies left or expanded to further enrich edited cells.

6. After drug selection, it usually less than 1 week to generate enough cells for genomic DNA extraction (> 5 X 10^5 cells). Before that, pooled edited cells can also be split for preliminary imaging verification. By using a 15-well imaging slide which is compatible with high-resolution microscopy (ibidi 81506), only a few cells are needed (3 X 10^3 cells/well) due to the small size of well.
7. Preliminary imaging verification can be used to check whether protein expression, subcellular localization, and dynamics are uniform and consistent across pooled cells.

### 6. Perform PCR genotyping in pooled cells (<1d)

**Aim:** After transfecting and enriching drug-resistant cells, we can pool putatively edited cells and verify insertions using PCR. A specific band should be observed in the pooled knock-in cells compared with Wild-type (WT) cells, indicating a successful insertion (Figure 9). PCR genotyping requires a forward primer upstream of the target sgRNA site and a reverse primer inside the minicircle donor. The forward primer is specific to the target gene, while the reverse primer is universal.

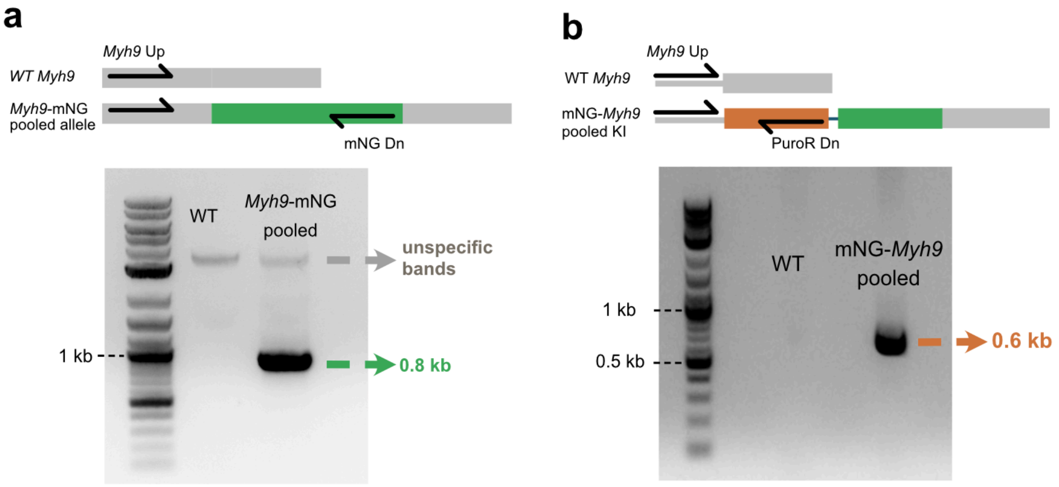

**Figure 9 PCR genotyping of pooled cells**
PCR genotyping requires a forward primer upstream of the sgRNA target site and a reverse primer inside the donor. **9a.** *Myh9* Up and mNG Dn primers amplified a specific band in pooled cells compared to WT cells, indicating a successful C-terminal insertion. **9b.** *Myh9* Up and PuroR Dn primers also gave rise to a specific band in pooled cells compared to WT cells, indicating a successful N-terminal insertion.

**Useful tools:**

1. NCBI/gene: https://www.ncbi.nlm.nih.gov/gene
2. CHOPCHOP: https://chopchop.cbu.uib.no/
3. Primer BLAST: https://www.ncbi.nlm.nih.gov/tools/primer-blast/

**Materials:**

1. PureLink Genomic DNA Mini Kit (Thermo Fisher K182001)
2. LongAmp Taq 2X Master Mix (NEB M0287S)

3. Primers: a forward primer upstream of the cutting site, and a reverse primer inside donor vector (Table 2).

| Tag | Resistance | NT or CT | Reverse primer for genotyping |
|---|---|---|---|
| mNG | PuroR | CT | mNG Dn: 5'-TTTGTAGAGCTCATCCATGC-3' |
| mNG | PuroR | NT | PuroR Dn: 5'-CATAGAAGGGAAGATTCCTTG-3' |
| Halo | HygroR | CT | Halo Dn: 5'-ACCGCTAATCTCCAAAGTT-3' |
| Halo | HygroR | NT | HygroR Dn: 5'-AAGCACTTCAACACAACCAT-3' |

**Table 2: Reverse primer for genotyping**
There are choices of different reverse primers depending on different tags and terminal insertions. Abbreviations: C-terminal (CT), N-terminal (NT), mNeonGreen (mNG), Puromycin resistance (PuroR), HaloTag (Halo), Hygromycin resistance (HygroR).

**Protocol:**

1. Search a target gene in NCBI/gene website, then click the correct search result based on organism and gene name. Scroll down to "Genomic regions, transcripts, and products" and go to nucleotide by opening "GenBank" directory. On the GenBank page, click "Send to", check "Complete Record" and "File", make sure "Format" is "GenBank", click "Create File" to download the WT genomic sequence of the target gene.
2. Identify the cut site in the downloaded genomic sequence. Cas9 cuts 3-nucleotides upstream of the PAM site. Then, insert (i.e. copy and paste) the sequence of linearized donor into the cut site to produce the sequence of the desired knock-in allele *in silico.*
3. Design a forward primer upstream of the cut site. Use Primer BLAST to design a forward primer because it can help to avoid any unspecific bands in the genotyping reaction.
4. Split pooled cells for genomic DNA extraction and subculture.
5. Take 5 X 10^5 cells of pooled edited and WT cells and extract genomic DNA using the PureLink Genomic DNA Mini Kit.
6. Set up PCR reactions using 50 ng of genomic DNA from pooled edited or WT cells as template, 1 µL of 10 µM forward primer and 1 µL of 10 µM reverse primer, 12.5 µL of LongAmp Taq 2X Master Mix, and Nuclease-free water to a final volume of 25 µL.
7. Run 30 cycles of PCR, following the instructions for LongAmp Taq.
8. Visualize the results on an agarose gel.

**Notes:**

1. 5 X 10^5 cells can yield around 6.25 µg of genomic DNA (250 ng/ µL in 25 µL).
2. The recommended amounts of genomic DNA template for LongAmp PCR are 1 ng–500 ng.
3. By using a pair of specific primers for the knock-in allele, only knock-in cells are expected to generate a specific band.
4. WT genomic DNA is required as a negative control to rule out non-specific bands.
5. Once pooled cells have been verified by PCR genotyping, the remaining cells can be used for isolation of clonal cell and cryopreservation of pooled cells.

### 7. Isolate monoclonal cells (1w)

**Aim:** To generate monoclonal cell lines, pooled edited cells can be seeded at low density so that cells are sparse and individual cells can generate single clonal colonies**.** When a single colony is big enough to be seen with a naked eye, it can be manually picked and expanded for further characterization.

**Materials:**

1. 10cm plate
2. 96-well U-bottom plates
3. Accutase (Sigma SCR005)
4. Growth medium 2i/LIF: Commercial N2B27(ESGRO Complete Basal Medium, Sigma SF002-500) or Homemade N2B27[6] complemented with 3 μM of GSK3β inhibitor (Sigma SML1046), 1 μM of MEK inhibitor (Sigma PZ0162) and 100 U/mL of Leukemia Inhibitory Factor (Sigma ESG1106)

**Protocol:**

1. On d0, seed 1 X 10^4 pooled edited cells with 10 mL of 2i/LIF on a gelatin-coated 10cm plate.
2. Change 2i/LIF every day until the colonies are big enough to see with a naked eye.
3. On d7, the colonies are typically big enough to be singled. Prepare two U-shape-bottom 96-well plates, one with 20 μL Accutase in each well, the other coated with gelatin. Pick individual colonies with a P10 pipet, set at 5 μL, under a dissection microscope. Transfer each colony into the Accutase-containing wells.
4. After picking the desired number of clones, incubate the Accutase-containing wells at 37°C for 10 min.
5. Add 150 μL of 2i/LIF to the Accutase plate containing colonies, pipet the mixture up and down to dissociate colonies into single cells, then transfer cell suspension to gelatin-coated plate.
6. On the next day, gently remove most of medium but leave enough to cover the cells, add fresh 2i/LIF.
7. Change 2i/LIF every day until cells reach confluence.

**Notes:**

1. To ensure isolation of true single cell clones, pick colonies with a similar size, and ensure that they are far enough away from the other colonies.
2. Single healthy-looking colonies (shiny and dome-shaped cell cluster with smooth and sharp borders) and change pipette tips between picks.
3. Microscopy can be helpful to verify that only one clone is picked to avoid none or more than one clone in one well.
4. Do not leave colonies in Accutase for more than 15-20 min.
5. Typically, 12 clones are sufficient to obtain several functional knock-in cell lines. If you need bi-allelic tagging (i.e., both genomic copies are tagged) or a seamless junction between the tag and the gene of interest, it may require isolation of more clones. It depends on the cutting efficiency of Cas9 towards the gene of interest.
6. A multi-channel pipet makes manipulation of multiple colonies in 96-well plates much more efficient.
7. Don't detach cells when changing medium. If you find cells can't attach to the bottom of the 96-well plates, 1% FCS in 2i/LIF can help cell adhesion.

### 10. Perform direct PCR genotyping in clonal cells (1-2w)

**Aim:** Quick lysis saves time and labor when genotyping multiple clones. To verify an insertion in clonal cells, PCR genotyping requires a forward primer upstream of the sgRNA target site. The reverse primer can be either downstream of the target site or inside the donor vector. To distinguish whether the insertion is mono-allelic (i.e., only one genomic copy is tagged) or bi-allelic, PCR genotyping requires a pair of primers upstream and downstream of the cutting site. For example, PCR with β-catenin Up and mNG Dn primers gave rise to a specific 0.8 kb band in knock-in clones (20.8 and 20.11), but no band in WT clones. PCR with β-catenin Up and Dn primers amplified a 0.3kb band in WT cells and for the non-edited allele. The knock-in allele produced a larger band of 1.8 kb (β-catenin 0.3 kb + minicircle donor 1.5 kb). The presence of both bands indicates a mono-allelic tag (e.g., clone 20.8), while the loss of short band indicates the possibility of a bi-allelic clone (e.g., clone 20.11). Sanger sequencing of gel-purified PCR products can be used to validate PCR results. Figure 10 shows an example of the expected results.

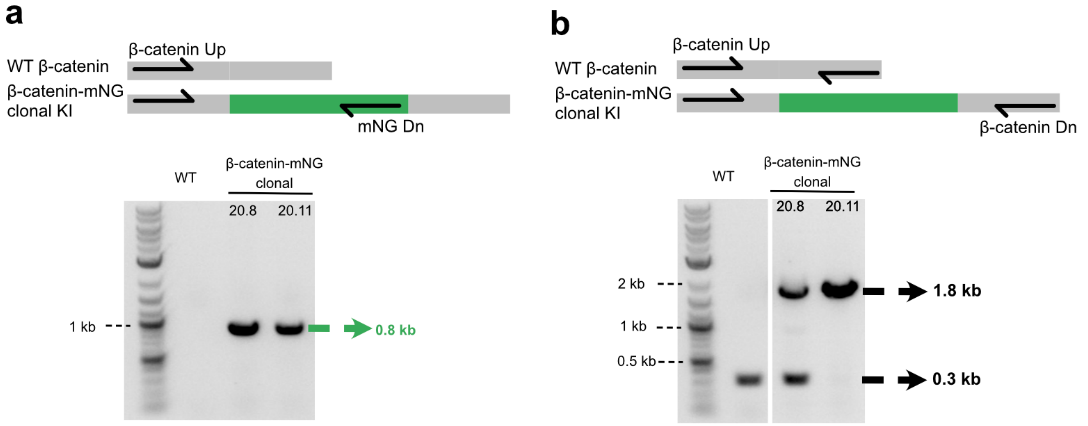

**Figure 10 PCR genotyping of clonal cells**
PCR genotyping requires a forward primer upstream of the sgRNA target site, and a reverse primer inside the donor or downstream of the target site. **10a.** β-catenin Up and mNG Dn primers amplified a specific band in clonal knock-in cells compared with WT cells, indicating a successful insertion. **10b.** β-catenin Up and Dn primers gave rise to both short and long bands in cell clone 20.8, suggesting mono-allelic tagging. The absence of a short band in clone 20.11 indicates a possible bi-allelic tag.

**Materials:**

1. Gelatin-coated 96-well and 24-well plates
2. Accutase (Sigma SCR005)
3. Growth medium 2i/LIF: Commercial N2B27(ESGRO Complete Basal Medium, Sigma SF002-500) or Homemade N2B27[6] complemented with 3 µM of GSK3β inhibitor (Sigma SML1046), 1 µM of MEK inhibitor (Sigma PZ0162) and 100 U/mL of Leukemia Inhibitory Factor (Sigma ESG1106)
4. Wash buffer: 500 mL of DMEM/F12 (Sigma D6421) + 8 mL of bovine serum albumin fraction V (Thermo Fisher 15260037)
5. Quick lysis buffer: 10 mg/mL proteinase K powder (Sigma P6556), 50 mM KCl, 10 mM Tris-HCl pH=8, 2 mM $MgCl_2$, 1% (v/v) NP40 (Thermo Fisher 85124), 0.45%(v/v) Tween 20 in milli-Q water.
6. PCR Tube Strips (VWR 89093-042)

7. LongAmp Taq 2X Master Mix (NEB # M0287)
8. Genotyping primers

**Protocol:**

1. When clonal cells reach confluence in the 96-well plate, split cells into two sets of plates, one in 96-well plate for PCR genotyping, the other in 24-well plate for subculture.
2. Change 2i/LIF every day until cells in 96-well plate reach confluence
3. Aspirate all the medium in 96-well plate, add 100 µL of Quick lysis buffer, pipet up and down to dissociate colonies, then transfer the cell lysates to PCR Tube Strips. Seal the plate and incubate at 65°C tor 2 h then heat at 95°C for 10 min.
4. Dilute cell lysates 1:20 with Nuclease-free water
9. Set up PCR reactions using 1 µL of diluted cell lysates or WT genomic DNA as template, 1 µL of 10 µM forward primer and 1 µL of 10 µM reverse primer, 12.5 µL of LongAmp Taq 2X Master Mix, and Nuclease-free water to 25 µL.
10. Run 30 cycles of PCR, following the instructions for LongAmp Taq.
11. Visualize the results on an agarose gel.

**Notes:**

1. The primers upstream and downstream of the cleavage site on genomic DNA are based on your chosen sgRNA. We design primers that center the cutting site within a region of 100-300bp.
2. Primer-BLAST is recommended to design primers for PCR genotyping to avoid non-specific bands.
3. Clones lacking inserts can safely be discarded at this stage.
4. Positive PCR products from step 9 can be gel purified and sent for Sanger sequencing to determine the nature of the insertion. Based on this information, decide which clones to keep. Typically, we maintain at least five different clonal cell lines for each modification to make sure the phenotype is consistent.

1. Ezkurdia, I. *et al.* Most Highly Expressed Protein-Coding Genes Have a Single Dominant Isoform. *J Proteome Res* **14**, 1880–1887 (2015).

2. Rodriguez, J. M. *et al.* APPRIS: annotation of principal and alternative splice isoforms. *Nucleic Acids Res* **41**, D110–D117 (2013).

3. Chung, S. *et al.* Analysis of Different Promoter Systems for Efficient Transgene Expression in Mouse Embryonic Stem Cell Lines. *Stem Cells* **20**, 139–145 (2002).

4. Dickinson, D. J. & Goldstein, B. CRISPR-Based Methods for Caenorhabditis elegans Genome Engineering. *Genetics* **202**, 885–901 (2016).

5. Kay, M. A., He, C.-Y. & Chen, Z.-Y. A robust system for production of minicircle DNA vectors. *Nat Biotechnol* **28**, 1287–1289 (2010).

6. Mulas, C. *et al.* Correction: Defined conditions for propagation and manipulation of mouse embryonic stem cells (doi:10.1242/dev.173146). *Development* **146**, dev178970 (2019).

7. Tamm, C., Kadekar, S., Pijuan-Galitó, S. & Annerén, C. Fast and Efficient Transfection of Mouse Embryonic Stem Cells Using Non-Viral Reagents. *Stem Cell Rev* **12**, 584–591 (2016).
